# A paleogenomic investigation of overharvest implications in an endemic wild reindeer subspecies

**DOI:** 10.1101/2023.09.21.558762

**Authors:** Fabian L. Kellner, Mathilde Le Moullec, Martin R. Ellegaard, Jørgen Rosvold, Bart Peeters, Hamish A. Burnett, Åshild Ønvik Pedersen, Jaelle C. Brealey, Nicolas Dussex, Vanessa C. Bieker, Brage B. Hansen, Michael D. Martin

## Abstract

Overharvest can severely reduce the abundance and distribution of a species and thereby impact its genetic diversity and threaten its future viability. Overharvest remains an ongoing issue for Arctic mammals, which due to climate change now also confront one of the fastest changing environments on Earth. The high-Arctic Svalbard reindeer (*Rangifer tarandus platyrhynchus)*, endemic to Svalbard, experienced a harvest-induced demographic bottleneck that occurred during the 17-20^th^ century. Here we investigate changes in genetic diversity, population structure and gene-specific differentiation during and after this overharvesting event. Using whole-genome shotgun sequencing, we generated the first ancient nuclear (*n* = 11) and mitochondrial (*n* = 18) genomes from Svalbard reindeer (up to 4000 BP) and integrated these data with a large collection of modern genome sequences (*n* = 90), to infer temporal changes. We show that hunting resulted in major genetic changes and restructuring in reindeer populations. Near-extirpation and 400 years of genetic drift have altered the allele frequencies of important genes contributing to diverse biological functions. Median heterozygosity was reduced by 23%, while the mitochondrial genetic diversity was reduced only to a limited extent, likely due to low pre-harvest diversity and a complex post-harvest recolonization process. Such genomic erosion and genetic isolation of populations due to past anthropogenic disturbance will likely play a major role in metapopulation dynamics (i.e., extirpation, recolonization) under further climate change. Our results from a high-arctic case study therefore emphasize the need to understand the long-term interplay of past, current, and future stressors in wildlife conservation.

## 1. Introduction

Excessive harvest reduces population size and genetic diversity and can ultimately lead to local extirpation or global (i.e., species) extinction (Frankham, 2005; Spielman et al., 2004). Overharvesting, also called overexploitation, has for long impacted fish and large mammals and occur today in combination with climate change and habitat loss (Bowyer et al., 2019; IUCN, 2020; Lorenzen et al., 2011; Luypaert et al., 2020). Populations may recover demographically after overharvest and near-extinction events, but their genetic diversity can remain low (Lande et al., 2003). Harvest-induced bottlenecks will likely also reduce a species’ resilience and adaptive capabilities when facing future challenges such as global climate change (Frankham et al., 2002). Nevertheless, despite that genetic diversity, along with species and ecosystem diversity, is recognized as one of the three pillars of biodiversity, it is not yet widely considered by conservation policymakers (Jensen et al., 2022; Laikre et al., 2010). It is therefore important to quantify the loss of genetic variation and changes in population genetic structure following such bottlenecks, in order to appropriately set conservation measures and to better predict the trajectory of the potential recovery.

Contemporary genetic material can hold considerable information about past demographic processes. For instance, methods employing coalescent theory can retrospectively detect genetic bottlenecks, and inform on their magnitude and timing (Drummond et al., 2005). However, information about lineages that went extinct during the bottleneck is lost. Therefore, investigation of effects of past bottlenecks like overharvesting from only contemporary material may overlook the severity of the event (Leonardi et al., 2017). Advances in the fields of genomics and ancient DNA (aDNA) enable population level whole genome sequencing of nuclear and mitochondrial genomes, for instance of specimens living prior to harvest-induced bottlenecks (Leonardi et al., 2017; Mitchell & Rawlence, 2021). Ancient DNA is therefore a powerful tool that allows for direct temporal comparisons, setting a “baseline” for the state of the species before anthropogenic intrusions (Díez-Del-Molino et al., 2018; Jensen et al., 2022). The respective measures of genetic change through time are valuable in ecological, evolutionary and conservation contexts, with a potential to inform future conservation efforts (Jensen et al., 2022), for example by studying genomic erosion (Robin et al., 2022; Sánchez-Barreiro et al., 2021), the sum of genetic threats to small populations, such as decreasing genome-wide diversity, increasing genetic load and inbreeding, and reduced genome wide heterozygosity (Díez-Del-Molino et al., 2018; Frankham, 2005; Kohn et al., 2006). By comparing the genomes of different temporal populations, regions of high genomic divergence can be identified. Genes within these regions likely experienced evolution due to selection or genetic drift (Allendorf & Hard, 2009; Therkildsen et al., 2019)

Because wildlife extirpations (i.e., local population extinctions) are expected to accelerate in the future (IUCN, 2020), knowledge of past genetic changes is crucial to predict the future population genetics of populations that are currently in decline due to harvesting, climate change, habitat loss, or competition with invasive species. Many Arctic mammals, such as bowhead whale (*Balaena mysticetus* Linnaeus, 1758) and walrus (*Odobenus rosmarus* (Linnaeus, 1758)), as well as some reindeer and caribou (*Rangifer tarandus* Linnaeus, 1758) subspecies, experienced large-scale local extirpations about a century ago (CAFF, 2013). Some of the species were even driven to extinction (Byun et al., 2002; Gravlund et al., 1998).

The wild Svalbard reindeer (*Rangifer tarandus platyrhynchus,* Vrolik, 1829), a subspecies endemic to the Svalbard archipelago with distinct morphological and behavioral characteristics, was hunted down to approximately 1000 individuals and extirpated from ∼60% of its range before being protected by law in 1925 (Lønø, 1959). The reindeer subspecies survived in four remote locations (populations), from which they then slowly recolonised the archipelago, partially assisted by two translocation programs (Aanes et al., 2000), to reach a current population size at ∼22,000 individuals (Le Moullec et al., 2019). Conveniently, the cold and dry Arctic environment physically preserves ancient skeletal material and its genetic information to a relatively high degree. Ancient DNA from bones and antlers from Svalbard reindeer that lived prior to the presence of humans (before the 17^th^ century) therefore represent a unique opportunity to quantify effects of harvest-induced bottlenecks on the genetic composition of present-day metapopulations (Le Moullec et al., 2019). Thus, by comparing contemporary DNA with ancient DNA from Svalbard reindeer this ‘natural experiment’ can provide valuable information on the genetic consequences of harvest-induced bottlenecks and population recovery following successful conservation efforts in large animals.

Reindeer most likely colonized Svalbard from Eurasia through intermediate colonization of the Franz Josef Land archipelago as a stepping-stone (Kvie et al., 2016). The earliest evidence of reindeer presence on Svalbard is from 5,000-3,800 BP (van der Knaap, 1989). The locations of a sample of carbon-dated ancient bones suggest they subsequently spread across the entire Svalbard archipelago (Le Moullec et al., 2019; van der Knaap, 1989). Harvest was introduced when Svalbard was discovered in 1596, but the most intensive hunting period of reindeer occurred in the early 1900s (Hoel, 1916; Lønø, 1959; Wollebaek, 1926), until protection in 1925. Lønø et al. (1959) documented that reindeer had then survived at low abundance in Nordenskiöld Land (central Spitsbergen), Reinsdyrflya (North Spitsbergen), Nordaustlandet (North East Svalbard) and Edgeøya (East Svalbard). Since then, reindeer have recolonised most of its former range from these four remnant populations (Peeters et al., 2020).

Six genetically distinct reindeer groups are now present on the Svalbard archipelago: four populations that expanded from their respective ‘hunting refugia’, and two populations founded by individuals from central Spitsbergen. Of the latter, one was founded along the west coast of Spitsbergen following two translocations, and another was founded by natural recolonization to south Spitsbergen, with strong genetic drift following gradual expansion (Burnett et al., 2022; Peeters et al., 2020). Svalbard reindeer disperse slowly due to their sedentary behavior and the fragmented landscape (Le Moullec et al., 2019). Habitat connectivity is further reduced with climate warming and declining cover of coastal sea-ice, which provides an important dispersal corridor (Peeters et al., 2020). Because hunting, predation, insect harassment and intra-specific competition for resources play only minor roles (Derocher et al., 2000; Reimers, 1984; Stempniewicz et al., 2021; Williamsen et al., 2019), population growth is mainly determined by the density-dependent weather effects on access to resources in winter (Albon et al., 2017; Hansen et al., 2019; Loe et al., 2020).

While some wild reindeer/caribou subspecies are undergoing strong declines due to anthropogenic landscape fragmentation and climate change (Collard et al., 2020; Festa-Bianchet et al., 2011), the Svalbard reindeer is increasing in abundance (Hansen et al., 2019; Le Moullec et al., 2019). This increase is mainly driven by recovery from the past overharvesting and by climate change improving and enhancing the length of snow-free season (Hansen et al., 2019; Le Moullec et al., 2019; Loe et al., 2020). The Svalbard reindeer genetic diversity is by far the lowest among the Rangifer subspecies (Kvie et al., 2016; Yannic et al., 2013). Despite this, local variation in genetic diversity is strong, with decreasing diversity from central Spitsbergen towards the peripheries of the archipelago (Peeters et al., 2020), (Burnett et al., 2022; Kvie et al., 2016). Inbreeding (i.e. long runs of homozygosity) is stronger in the non-admixed naturally recolonized populations than in the two translocated populations, likely as a result of having experienced a series of bottlenecks (Burnett et al., 2022). The translocated populations largely maintained the genetic diversity level of their source populations, despite <15 founder individuals for each translocation, likely because of rapid population growth and overlapping generations (Burnett et al., 2022).

Here, we investigate the genetic impacts of the population bottleneck caused by the past overharvest of Svalbard reindeer. We hypothesize that the reindeer suffered genomic erosion due to this overexploitation event. To infer changes in genetic structure, diversity and allele frequencies, we integrated paleogenomic data with a large dataset of contemporary genome sequences and estimated genetic diversity and genomic erosion before (4,000-400 calibrated years Before Present [BP]), during (500–0 BP, equating to 1500–1950 Common era [CE], also corresponding to the period with uncertain carbon-dating) and after the overharvesting period (> 1950 CE).

## 2. Materials and Methods

### 2.1 Study system and sample acquisition

The Svalbard archipelago (76°–81°N, 10°–35°E) is surrounded by the Greenland and Barents Seas, south of the Arctic Ocean. The archipelago consists of over 500 islands, the largest being Spitsbergen, Nordaustlandet, Edgeøya, Barentsøya and Prins Karl Forland. Reindeer inhabit vegetated land patches, which make up only 16% of the land cover, fragmented by fjords and tide-water glaciers (Johansen et al., 2012). Central Spitsbergen and Edgeøya holds a network of unglaciated valleys with the highest density of reindeer (Le Moullec et al., 2019).

At the time of the oldest Svalbard reindeer records (∼5,000 BP), the Svalbard summer climate was approximately 1.5-2°C warmer than the recent reference period of 1912-2012 CE (van der Bilt et al., 2019). The climate became progressively cooler over time until the pre-industrial period at around 1500 CE (van der Bilt et al., 2019), with increasing sea-ice cover in the Fram Strait (Werner et al., 2016). Currently, Svalbard is among the regions on Earth experiencing the strongest temperature increase, with 0.5°C increase per decade in summer and 1.3°C increase per decade for year-round measurements at Svalbard Airport, 2001-2020 CE, (Isaksen et al., 2022). Partly related to this, the sea-ice concentration around Svalbard has decreased drastically in recent years at a rate of 10-15% per decade in winter (2001-2020 CE, (Isaksen et al., 2022). The West side of Spitsbergen is now ice-free year-round, except for some inner fjords or in front of some tide-water glaciers.

Subfossil bones and antlers (*n* = 18) were collected from various sites across the Svalbard archipelago during 2014-2015 CE as described previously (Le Moullec et al., 2019). Subfossil collection was approved by the Governor of Svalbard (RIS-ID: 10015 and 10128). Sample ages were determined via ^14^C dating (Table 1). All ^14^C dates were calibrated with the IntCal20 calibration curve (Reimer, 2020) using the *calibrate* function in the package *rcarbon* (Crema & Bevan, 2021) in R v4.1.0 (R Core Team, 2021). The subfossil materials were combined with a previously published genomic dataset from samples collected in 2014-2018 and consisting of 90 contemporary Svalbard reindeer from a recent population genomics study (Burnett et al., 2022) PRJEB57293(Burnett et al., 2022). To put harvest-induced changes within Svalbard reindeer populations into a temporal context, we subdivided the samples into three time periods: Before (4,000-400 years before present, BP, hereafter referred to as *pre-hunting*), during (400 years BP-1950 Common Era, CE, hereafter referred to as *during-hunting*), and after (> 1950 CE, hereafter referred to as *post-hunting*) the major harvest-induced bottleneck that occurred from the 17th to the early 20th century (Lønø 1959).

**Table 1.**
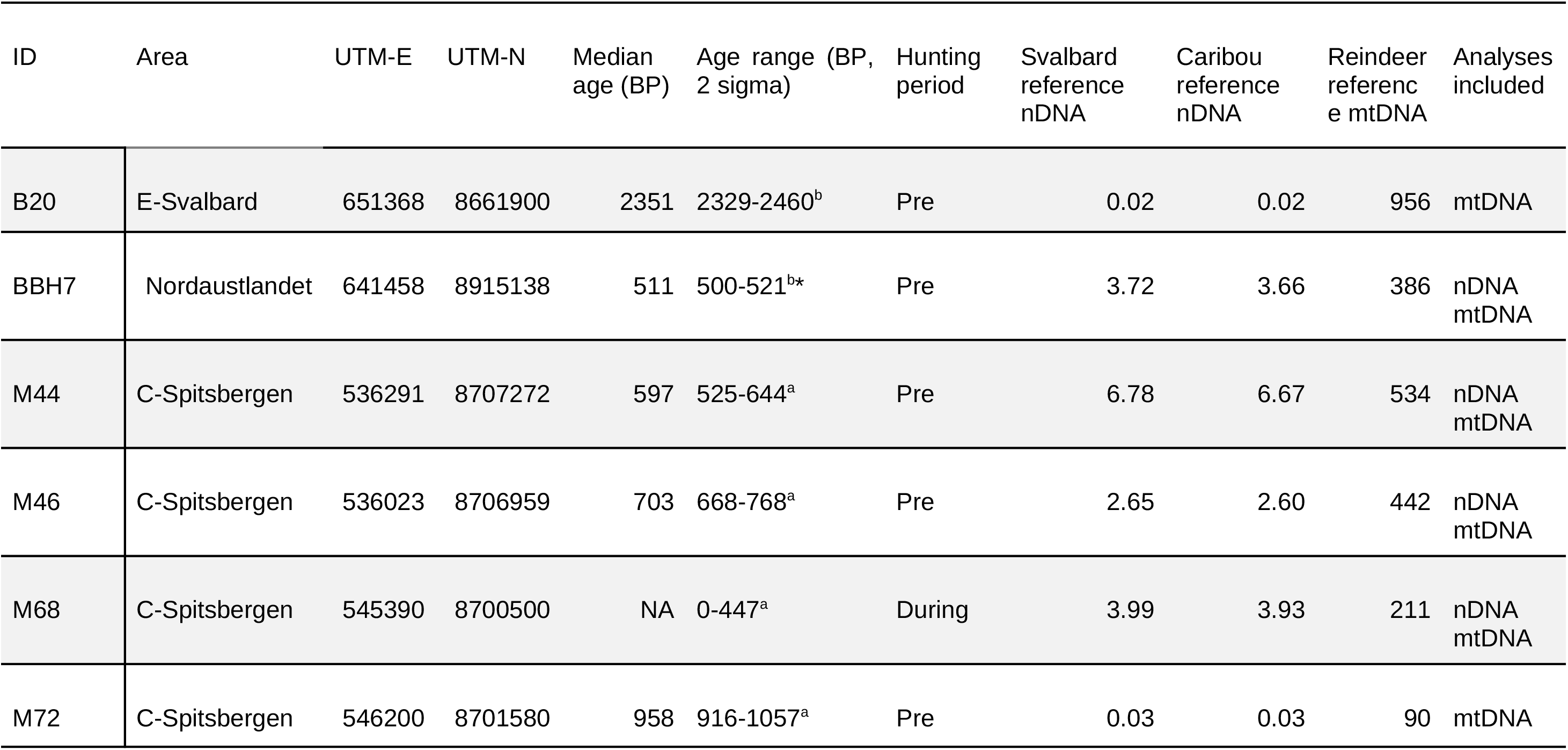
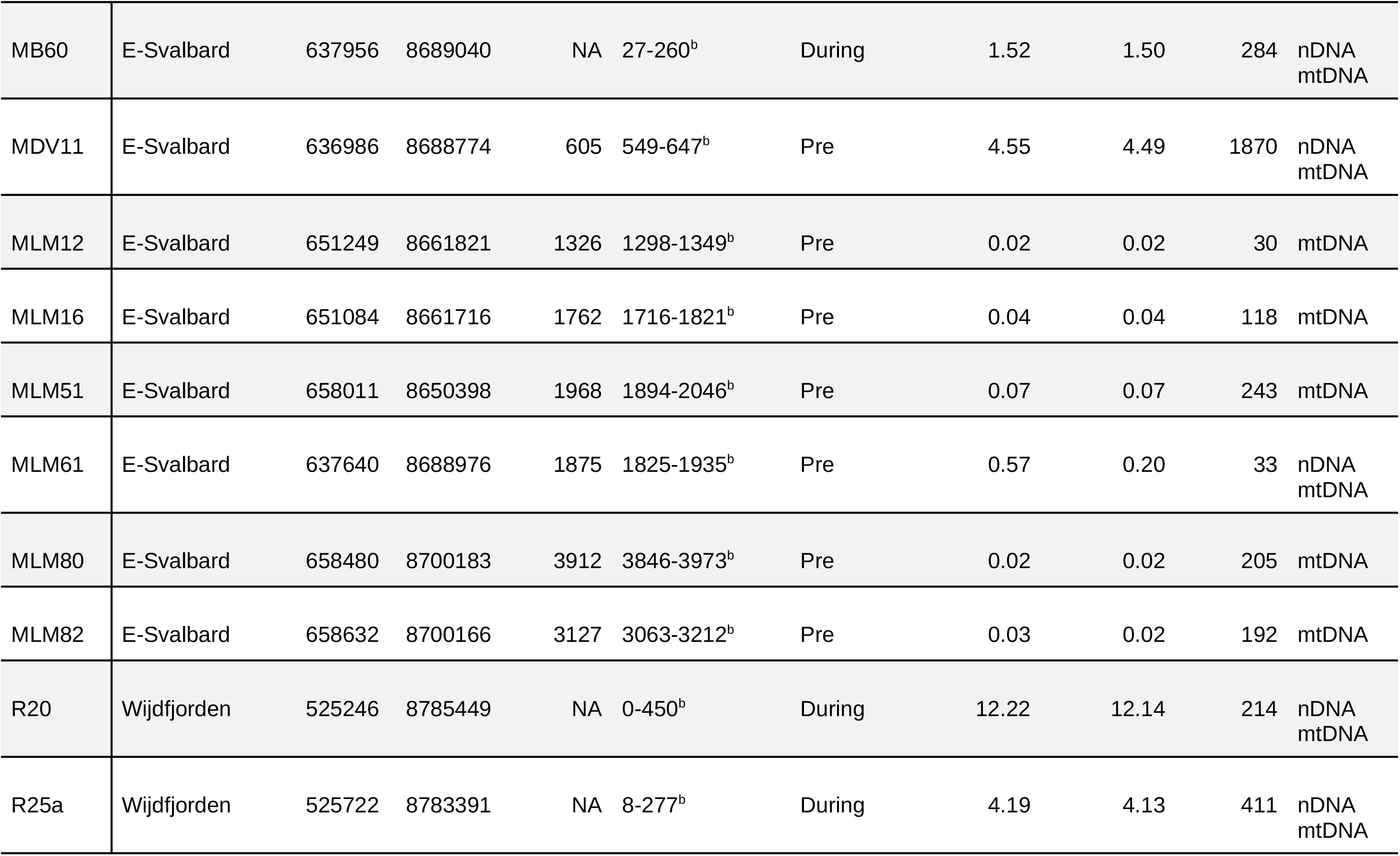
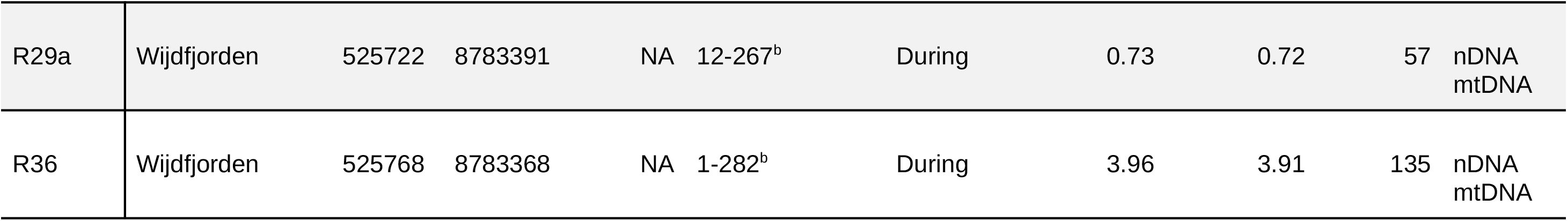
Ancient Svalbard reindeer sample overview. Sample provenance (‘E’ = East, ‘C’ = central) indicated with their UTM East and UTM North coordinates. Reported ages are calibrated radiocarbon-dated ages (IntCal20) in calendar years before present (BP, before 1950 CE). Summary of sample and mean mapping depths of ancient nuclear genomes (nDNA) against the Svalbard reindeer and caribou reference genome, as well as the reindeer mitochondrial (mtDNA) reference genome. Sample provenance, sequencing, and mapping statistics of ancient samples against the Svalbard reindeer reference genome. Reported ages are calibrated carbon-dated ages (IntCal20) in calendar years before present (BP, before 1950 CE). * = Radiocarbon date from previous study (Le Moullec et al., 2019). ^a^ = dated at the Uppsala Angström Laboratory. ^b^ = dated at the Norwegian University of Science and Technology, The National Laboratory of Ange Determination.

### 2.2 DNA extraction, library building, and sequencing

All pre-PCR manipulations of sample genetic material were conducted in a dedicated, positively pressurized ancient DNA laboratory facility at the NTNU University Museum. 48- 360 mg of bone material were collected using a Dremel disc drill and subsequently crushed to fragments of maximum 1-mm diameter. DNA was extracted with a custom, silica-based extraction protocol. For digestion, a custom digestion buffer consisting of 1.25% (v/v) proteinase K (20 mg/mL), 90% (v/v) EDTA (0.5M) and 8.75% (v/v) molecular-grade water was used. Samples were pre-digested in 1 mL digestion buffer for 10 minutes at 37°C on a rotor. The samples were spun down, the pre-digest was removed, and 4 mL digestion buffer was added to the samples. Samples were left for digestion on a rotor for 18 hours at 37°C. The lysis buffer was mixed 1:10 with Qiagen PB buffer modified by adding 9 mL sodium acetate (5 M) and 2 mL NaCl (5 M) to a 500 mL stock solution. pH was adjusted to 4.0 using concentrated (37% / 12M) HCl. 50 µL in-solution silica beads were added, and samples were left on a rotor for 1 hr at ambient temperature to allow binding of the DNA. Silica pellets with bound DNA were purified using Qiagen MinElute purification kit following the manufacturer’s instructions and eluted in 65 µL Qiagen EB buffer. For every extraction batch, an extraction blank was carried out alongside the samples.

For each sample and the extraction blanks, 32 μL of DNA extract were built into double-L of DNA extract were built into double- stranded libraries. Libraries were prepared following the BEST 2.0 double-stranded library protocol (Carøe et al., 2018). Three DNA extraction blanks were included, as well as a single library blank (no-template control). 10 μL of DNA extract were built into double-L of each library were amplified in 50 μL of DNA extract were built into double-L reactions with AmpliTaq Gold polymerase, using between 13 and 22 cycles and a dual indexing approach (Kircher et al., 2012). The optimal number of PCR cycles for each library was determined via qPCR on a QuantStudio3 instrument (ThermoFisher). The indexed libraries were purified using SPRI beads (Rohland & Reich, 2012) and eluted in 30 µL EBT buffer. The amplified libraries were quantified on 4200 TapeStation instrument (Agilent Technologies) using High Sensitivity D1000 ScreenTapes. The libraries were then pooled in equimolar concentrations and initially sequenced on the Illumina MiniSeq platform in order to quantify their complexity and endogenous DNA content. The libraries were subsequently subjected to several rounds of PE 150 bp sequencing on the Illumina NovaSeq 6000 platform (Norwegian National Sequencing Centre and Novogene UK).

### 2.3 Bioinformatic analysis and post-mortem DNA damage assessment

Sequence data from the pooled libraries were demultiplexed according to their unique P5/P7 index barcode combinations. Prior to mapping, residual adapter sequences were removed and reads shorter than 25 bases were discarded with AdapterRemoval v2.2.4. Raw reads were mapped initially against the caribou (*Rangifer tarandus caribou*) nuclear genome (Taylor et al., 2019) and the reindeer (*Rangifer tarandus tarandus*) mitochondrial genome (Z. Li et al., 2017) in the framework of the PALEOMIX pipeline v1.2.13.2 (Schubert et al., 2014) using the ‘mem’ algorithm of the Burrows-Wheeler Aligner (BWA) v0.7.16a (H. Li, 2013) and no mapping quality (MAPQ) score filtering. PCR duplicates were marked using picardtools v2.20.2 (“Picard Toolkit,” 2019) tool MarkDuplicates. Mapped reads were realigned with the Genome Analysis Tool Kit (GATK v3.8-0) indel realigner (McKenna et al., 2010). Following mapping, soft-clipped reads were removed with samtools v1.12 (H. Li et al., 2009). Then mapDamage v2.0.9 (Jónsson et al., 2013) was used to assess and plot ancient DNA damage patterns and to rescale base quality scores accordingly. Sequencing depth statistics were estimated with samtools depths at a minimum phred-scaled mapping quality of 30 (Table S1). All samples were sequenced to a minimum of 0.2X sequencing depth of the caribou nuclear genome assembly after read filtering.

### 2.4 Construction of a consensus Svalbard reindeer reference genome

In order to improve mapping rate and more accurately reflect the divergent genome of the Svalbard reindeer, a reference Svalbard reindeer nuclear genome was generated from the deepest sequenced contemporary individuals of each metapopulation (T-15, C7 and B2, see Table S1). BAM files were downsampled to 26.9X, the lowest depth among the three genomes, with the samtools v1.12 view command using the flags *-b* and *-s*. Individual bam files were combined into a single bam file. The new reference sequence was determined with angsd v0.931 (Korneliussen et al., 2014) using the *-doFasta 2* tool with the options *- doCounts 1*, *-explode 1*, *-setMinDepthInd 2*, *-remove_bads 1*, requiring a minimum read mapping quality of 30 and a minimum base quality of 20. All samples were mapped against this new reference genome using the same methods described above.

### 2.5 Determination of ancestral states

Ancestral states were inferred by mapping publicly available sequencing data from three closely related (Heckeberg & Wörheide, 2019) species (moose, *Alces alces*, bioproj: PRJEB40679 (Dussex et al., 2020); red deer, *Cervus elaphus*, bioproj: PRJNA324173 (Bana et al., 2018); white-tailed deer, *Odocoileus virginianus*, NCBI PRJNA420098; Accession No. JAAVWD000000000) against the caribou reference genome (Taylor et al., 2019). BAM files were downsampled to 12.3X, the lowest depth among the three genomes, with samtools v1.12 and merged into a single BAM file for all three species. The ancestral state was inferred by choosing the most common base at each site with angsd v0.931 using the *- doFasta 2* tool with the options *-doCounts 1*, *-explode 1*, *-remove_bads 1*, and *-uniqueOnly 1*, requiring a minimum read mapping quality of 30 and a minimum base quality of 20.

### 2.6 Selection of nuclear genomic loci for further analysis

Genomic positions suitable for further downstream analysis were computed for ancient and modern samples with the GATK v3.8-0 CallableLoci tool, requiring a minimum read mapping quality of 30 and a minimum base quality of 20. The input BAM files for the ancient individuals were generated by merging all of that group’s BAM files with samtools merge and replacing the read groups with the picard-tools v2.20.2 AddOrReplaceReadGroups tool. For modern samples, only a single BAM file (sample T-15) was used. The average depth of these merged BAM files was calculated with samtools depth at mapping quality 30 and base quality 20 (ancient = 43.6, modern = 57.4) . The minimum depth was calculated as one third of the mean depth (ancient = 15, modern = 20) and the maximum depth as twice the mean depth (ancient = 87, modern = 115). The files were then further processed with bedtools v2.30.0 (Quinlan, 2014) in order to be used with the *-sites* and *-rf* options of angsd v0.931. Sites that were marked with excessive coverage or poor mapping quality were excluded from downstream analysis.

### 2.7 Mitogenome alignment and haplotype analysis

Variants in the mitogenome were identified with GATK v4.2.5.0 using a reindeer mitochondrial genome sequence (Z. Li et al., 2017) as the reference. For this, haplotypes were called with GATK HaplotypeCaller requiring a minimum read mapping quality of 30. Variants were only called when they were above a minimum phred-scaled confidence threshold of 30 (*-stand-call-conf 30*). We verified that all samples’ mitochondrial genomes were sequenced to a minimum depth of 30x after removing reads with mapping quality below 30 (see Table 1). GVCF files were merged using the GATK GenomicsDBImport tool, and a joined SNP call was performed with GATK GenotypeGVCFs with the setting *-stand- call-conf 30.* Mitochondrial haplotypes in the output variant call format (VCF) file were used to populate a FASTA multiple sequence alignment file using a custom python2 script. In this alignment, insertions and deletions were ignored, and individual haplotypes were deemed ambiguous (‘N’) if the sequencing depth was less than 10 reads or if the genotype quality score was less than 20. Despite these measures, one sample (MB16 with 17% ‘N’) was ambiguously assigned an haplotype and therefore removed from the analysis.

We used the *ape* package (Paradis & Schliep, 2019) in *R* to import the 16,362-bp mitogenome sequence alignment and then extracted haplotypes with the *haplotype* function from the *pegas package* (Paradis, 2010). We visualized haplotype diversity with a network linking haplotypes based on a parsimony criterion minimizing the number of sites segregating between haplotypes (i.e., TCS algorithm). Such an algorithm uses an infinite site model to calculate a pairwise distance matrix of the haplotypes, with pairwise deletion of missing data. With the haploNet function (Templeton et al., 1992) from *pegas*, we plotted haplotype networks from individuals living before, during, and after the hunting period. Across all mitogenome sequences and within each time period, we calculated genetic diversity statistics in *pegas*, using the *haplo.div* function to calculate haplotype diversity (Nei & Tajima, 1981) and the *nuc.div* function to calculate nucleotide diversity (Nei, 1987). The number of segregating sites were summed from the *seg.sites* function in *ape*.

### 2.8 Genotype likelihood estimation

Genotype likelihoods for nuclear genome loci were estimated with angsd v0.931, excluding reads with multiple matches to the reference genome (option *-uniqueOnly 1*). Allele frequencies were estimated with the option *-doGlf 2* using the combined estimators for fixed major and minor allele frequency as well as fixed major and unknown minor allele frequency (*-doMaf 3*). Major and minor allele frequencies were inferred from genotype likelihood data (*- doMajorMinor 1*). Variant sites were considered when they had a minimum minor allele frequency of 5% and minimum number of informative individuals set to half the total number of individuals (*-minInd 50*). Reads with a phred-scaled mapping quality score below 30 were excluded, as were bases with a quality score below 20 as well as reads with a samtools flag above 255 (not primary, failure and duplicate reads) with the option *-remove_bads 1.* The first five bases were trimmed from both ends of reads (*-trim 5),* and the frequencies of the bases were recorded with *-doCounts 1.* The depth of each individual (*-dumpCounts 2*) and the distribution of sequencing depths (-doDepth 1) were recorded. Files to be used in subsequent analyses with Plink were generated (*-doPlink 2*). Genotypes were encoded (*- doGeno 2*) for downstream analysis. Posterior genotype probabilities were estimated based on allele frequencies as prior (*-doPost 1*). Genotypes were only considered if their posterior probability was above 95% (*-postCutoff 0.95)*. Genotypes were considered missing in cases when the individual depth was below 2 (*-geno_minDepth 2*). Before further analysis, sites in strong linkage disequilibrium were pruned with Plink v1.90 beta 6.24 (Chang et al., 2015) using a window size of 50 kbp, a step size of 3 kbp, and a pairwise r^2^ threshold of 0.5 (-*- indep-pairwise 50 3 0.5).* Subsequently, regions with low mapping quality, excessive coverage as identified with CallableLoci (see above), or associated with sex chromosomes (for methods see (Burnett et al., 2022)) were removed via a custom python script. Also excluded were sites that are not variant in both the ancient ancient and modern datasets.

### 2.9 Principal components and admixture analyses

We visually identified related clusters with covariance matrices for principal component analysis (PCA) using PCAngsd v1.10 (Meisner & Albrechtsen, 2018), running 10,000 iterations to ensure convergence. Samples were grouped both spatially and by time period (pre-, during-, and post-hunting). A spatial group was defined as all individuals from each time period that were sampled within 80 km around centroid points calculated by hierarchical clustering with the *Geosphere* package in R (see map in Figure 2 and Figures S7 - S15). Genetic structure was estimated with NGSadmix (Skotte et al., 2013), including only variant sites with a minimum minor allele frequency of 5% and minimum number of informative individuals set to half the total number of individuals (*-minInd 50*). The number of estimated ancestral populations *K* ranged from 2 to 10. For each value of *K*, each of ten replicates were run with different random starting seeds, and the replicate with the highest likelihood was used for plotting. An optimal value for *K* was estimated with the delta-*K* method (Evanno et al., 2005). Admixture diversity scores where calculated based on *K* = 5 using the R package *entropy,* as described in (Harismendy et al., 2019). In order to evaluate possible sampling bias we repeated the analysis with a reduced dataset (see below).

### 2.10 Estimation of heterozygosity

Site allele frequency likelihoods were calculated for each sample individually with angsd (command-line option *-doSaf 1*) with a minimum base quality of 20 and a minimum read mapping quality of 30, using only selected sites and regions described previously and removing transitions (command-line option *-noTrans 1*) as well as 5 bp from the ends of all reads (*-trim 5*) to reduce artifacts of ancient DNA damage. Reads with multiple best hits and non-primary, failed, and duplicate reads were removed from the analysis (command-line options *-uniqueOnly 1* and *-remove_bads 1*). To polarize the site allele frequency (SAF) likelihoods, a FASTA format file with ancestral states (see above) was supplied (command- line option *-anc*). The GATK model (command-line option *-GL 2*) was used to estimate genotype likelihoods. Then, the site frequency spectrum (SFS) was estimated with angsd realSFS based on the polarized SAF likelihoods. Heterozygosity was then calculated in R as described in the *angsd* documentation. A pairwise Mann-Whitney U test with default settings as implemented in R was performed to judge the significance of differences in heterozygosity between groups. We obtained the delta estimator (Díez-Del-Molino et al., 2018) of heterozygosity (*ΔHH*) by calculating *ΔHH* =(*med* (*H* 2) *−med* (*H* 1))/*med* (*H* 1), with *H_2_* being the set of individual genome-wide heterozygosity values in the younger sample set, *H_1_* being the set of individual genome-wide heterozygosity values in the older sample set, and med being the median value.

### 2.11 Temporal genomic differentiation

To assess the potential phenotypic impacts of overharvest on Svalbard reindeer, we identified genomic regions that were highly differentiated in a comparison of the three time groups among each other. To reduce potential biases introduced by the large size of the contemporary reindeer population sample, a smaller dataset of modern individuals representing all geographic regions was chosen semi-randomly (*n*=11). The selected samples were T-15, T-7 (Wijdefjorden), B1, B2, T-44 (Central-Spitsbergen), C6, C7 (East Svalbard), T-20 (West-Spitsbergen), C28 (Nordaustlandet), B-132, C-84 (South- Spitsbergen). Within-population site frequency spectra were calculated with angsd v0.931 using the options *-dosaf 1*, *-gl 1*, *-noTrans 1*, *-trim 5*, *-remove_bads 1*, *-minMapQ 30*, and *- minQ 20*. As described above, the SAF was polarized with ancestral states, and only selected sites and regions were used. The pairwise 2D-SFS was estimated with the angsd *realSFS* tool (Nielsen et al., 2012) and between-population differentiation *F_ST_* values were estimated with realSFS commands *fst index* and *fst stats*.

Additionally, an *F_ST_* sliding window analysis was performed as pairwise comparisons among the time groups, using the angsd *realSFS* command *fst stats2* with non-overlapping 10-kbp windows. *F_ST_* values were z-transformed around their mean, and windows with z ≥ 6 were defined as outlier windows. The neutrality test statistics Watterson’s theta (Watterson, 1975) and Fay and Wu’s *H* (Fay & Wu, 2000) were calculated within the same windows by using the angsd realSFS commands *saf2theta* and *thetaStat do_stat*. The mean per-site Watterson’s theta estimator was used to calculate effective population size (*N_e_*) assuming the average mammalian mutation rate of 2.2 ✕ 10^-9^ per site per year (Kumar & Subramanian, 2002) and a generation time of 6 years as previously estimated for Svalbard reindeer (Flagstad et al., 2022). Mean nucleotide diversity was calculated by dividing pairwise theta by the number of sites in each 10-kbp window.

We further investigated the roles and functions of genes within regions of high divergence between the during-hunting and post-hunting periods, which we defined as regions with mean *F_ST_* ≥ 0.5. We retrieved the amino acid sequences of known caribou genes from the annotation provided by (Taylor et al., 2019). Amino acid sequences of all 20,014 *Bos taurus* proteins were retrieved from UniProt (UniProt Consortium, 2021) and used to construct a blast protein database with the *makeblastdb* tool within blast+ v2.6.0 (Altschul et al., 1990). The sequences were identified via a *blastp* search against the *Bos taurus* protein database, only considering results with an e-value < 0.001. To select the best matching protein for each query sequence, the result with the smallest e-value was chosen. Ties were resolved by first considering highest bit-score, then percentage identity, and finally alignment length. We then identified which protein coding genes intersect with regions of high divergence.

## 3. Results

### 3.1 Read mapping and post-mortem DNA damage assessment

The final dataset consists of 12 individuals with calibrated median age ranging between 3,973–500 BP (hereafter referred to as the “pre-hunting” population), six individuals with age ranging between 447–0 BP (equating to 1503–1950 CE; hereafter referred to as the “during- hunting” population), and 90 present-day individuals (hereafter referred to as the “post-hunting” population). For genetic comparisons between DNA from subfossil bones/antlers and contemporary samples, the 18 pre-hunting and during-hunting individuals are collectively referred to as the “ancient” population, and the 90 post-hunting individuals are referred to as the “modern” population. For the nuclear genome analysis, seven pre-hunting samples were excluded due to their low sequencing depth, resulting in 11 ancient samples.

Nuclear genome raw mapping results are reported in Table S1. After filtering soft-clipped reads and reads with low mapping quality, the mean sequencing depth across all samples (including those that were only used for haplotype network analysis) mapped against the caribou reference assembly was 5.2X (mean ancient DNA: 2.3X; mean modern DNA: 5.7X) (see Table S2). The mean sequencing depth across all samples was slightly higher when mapping against the Svalbard reference genome (mean across all samples: 5.2X; mean ancient DNA: 2.4X; mean modern DNA: 8.8X). The mean sequencing depth of the mitochondrial genome across all samples was 1287.0X (mean ancient DNA: 340.2X; mean modern DNA: 1484.0X). The ancient genomes were confirmed to show characteristic ancient DNA damage patterns (see Figure S5).

### 3.2 Population structure

In order to assess population structure before, during and after the period of intense hunting, that resulted in local extirpation followed by recolonisation, we performed a PCA of nuclear genome variation and an analysis of genetic admixture based on nuclear genome genotype likelihoods, as well as constructed a haplotype network from mitogenome sequences. To identify patterns in these results, we then analyzed them according to the division of Svalbard reindeer metapopulation into 12 spatio-temporal groups.

After removing low-quality sites, 2,063,517,637 nuclear genomic positions were retained for analysis. Except for a few outliers, the post-hunting genomes cluster based on their geography in the PCA when considering the first two components PC1 and PC2 (Figure 1). There was a clear separation of eastern (East-Svalbard), northern (Wijdefjorden and Nordaustlandet) and southwestern (Central-, South- and West-Spitsbergen) individuals. Post-hunting individuals from southern, western, southwestern, and central Spitsbergen form a single tight cluster. The northern groups Nordaustlandet and Wiljdefjorden are not as clearly separated from each other, where the Wijdefjorden genomes form a particularly diffuse cluster. The ancient genomes, irrespective of their geographic assignment, form a loose cluster around the center of the PC2 axis and towards the left of the PC1 axis between the Nordaustlandet, Wijdefjorden and East-Svalbard clusters of post-hunting genomes.

**Figure 1:**
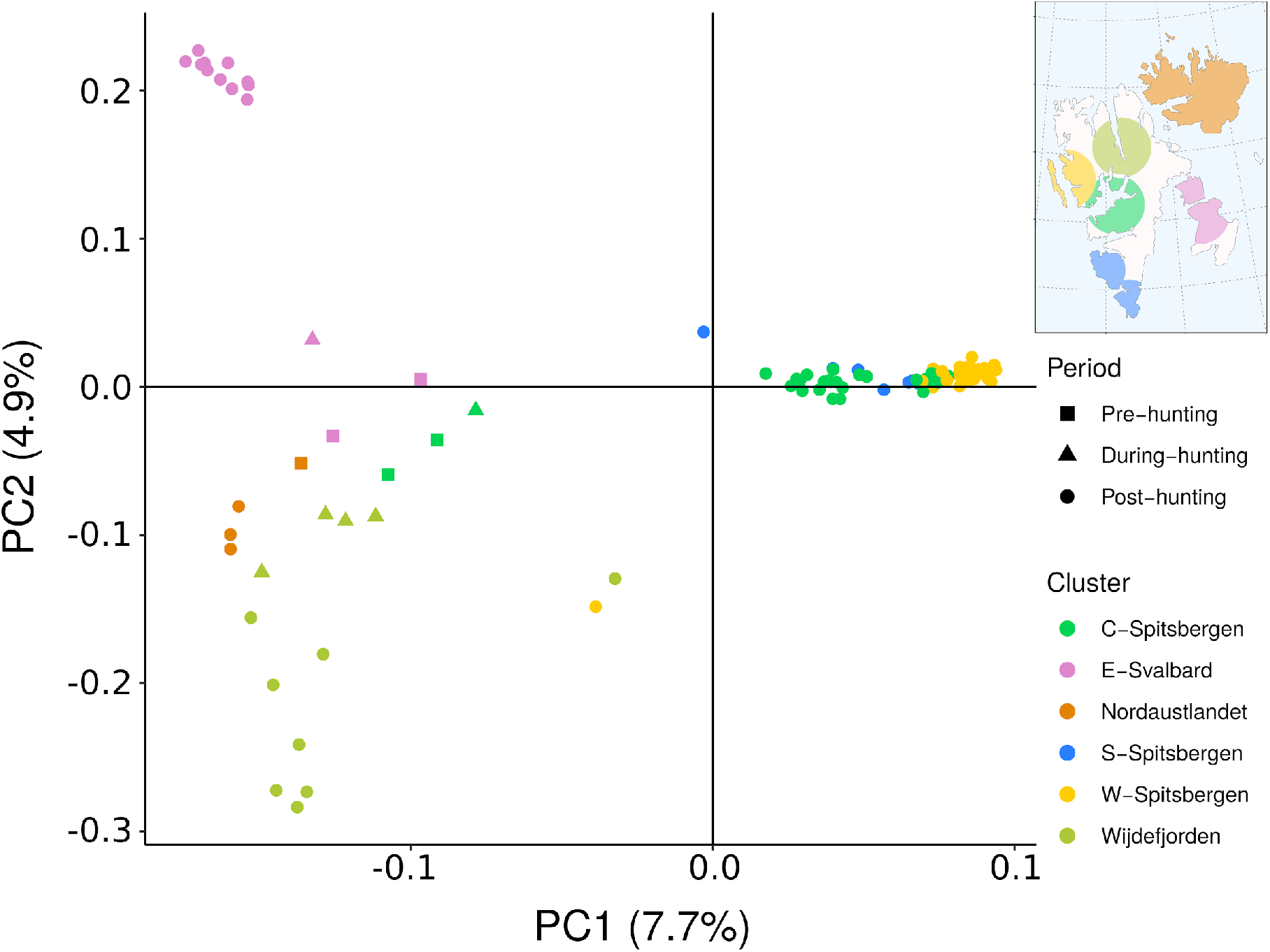
Principal component analysis (PCA) of the genomic variation of Svalbard reindeer. The first two principal components (PCs) are shown. Color denotes affiliation with a geographic cluster. Shape denotes affiliation with a time period. C-Spitsbergen = Central Spitsbergen; S-Spitsbergen = South Spitsbergen; W-Spitsbergen = West Spitsbergen; E- Svalbard = East Svalbard.

We performed an admixture analysis with a dataset including all individual nuclear genomes (Figure 2, Figures S6 and S16). The highest Δ*K* value was found for *K*=5 (2769.4, Fig. S17). The assignment of modern groups to ancestry clusters strongly correlates with geography. These five major genetic clusters of Svalbard reindeer are: a pink cluster, composed exclusively of post-hunting east Svalbard individuals (diversity score [DS] = 0); a blue cluster, characteristic for south Spitsbergen (DS = 0.151); a green cluster which is the main component of Wijdefjorden (DS = 0.402) individuals; an orange cluster characteristic of Nordaustlandet (DS = 0); and a yellow cluster that is the main component of central and west Spitsbergen (DS = 0.329). The genomes of individuals belonging to each geographic group in the more isolated outer regions of the archipelago are almost fully assigned to their own private ancestral population clusters (DS = 0 - 0.151), while the geographic group at the center of the archipelago (Central-Spitsbergen. DS = 0.640) shows admixture of genetic clusters dominant in southern, western, and northern populations (Figure 2). As expected, there is shared ancestry between Central-Spitsbergen and West-Spitsbergen, a population that was reintroduced from Central-Spitsbergen (Aanes et al., 2000). Only very little ancestry is shared between East-Svalbard and the other groups, and the same is true for Nordaustlandet. Overall, the ancestry assignments agree well with the groupings in the PCA.

**Figure 2:**
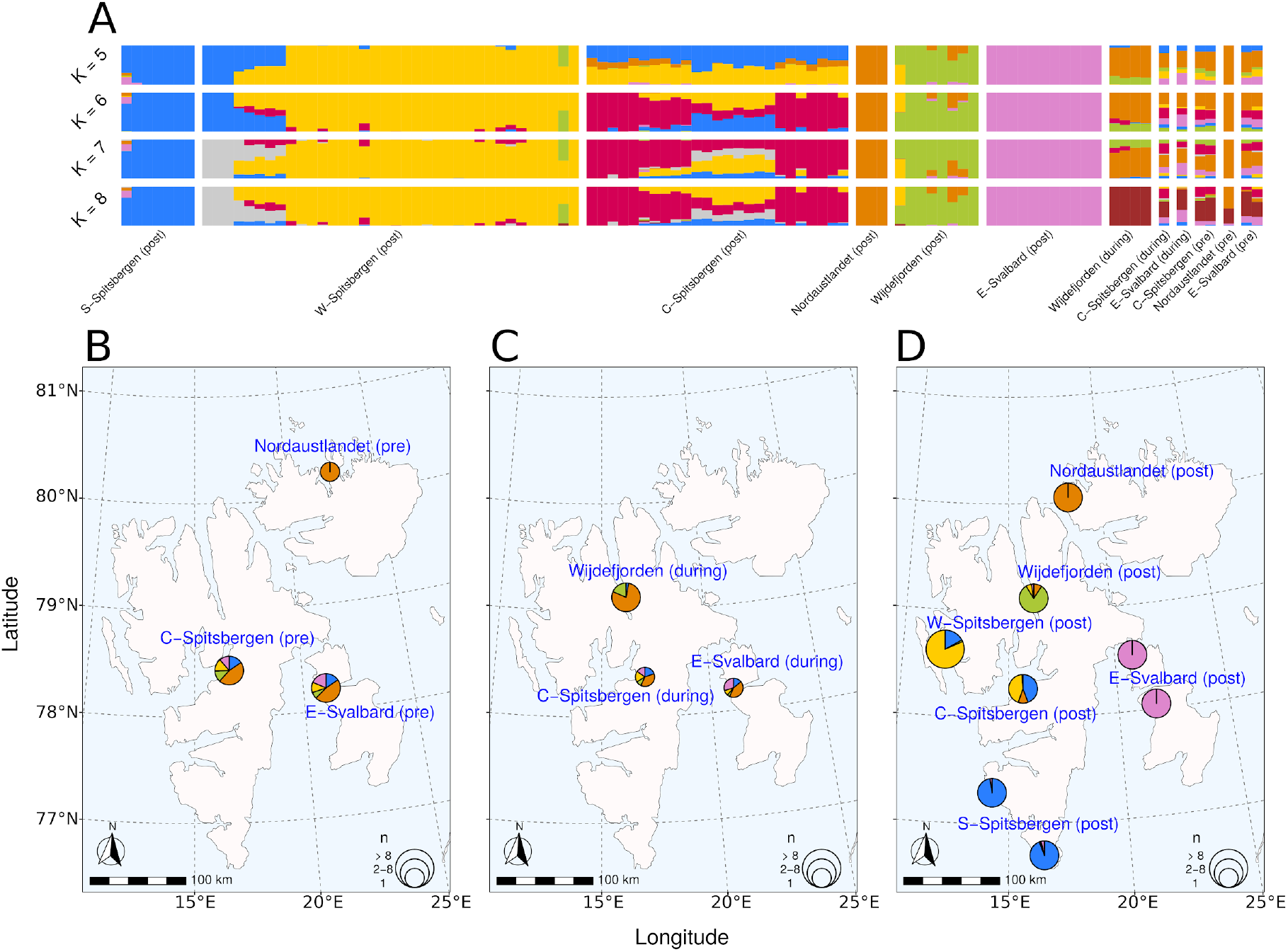
Admixture analysis of Svalbard reindeer in relation to sampling location. **A:** Bar plot of admixture proportions for *K* = 2 to *K =* 8. Each bar represents one individual Svalbard reindeer and the color represents affiliation to a proposed ancestral population. Individuals are grouped by spatiotemporal groups as defined by cluster analysis. **B-D:** Admixture proportions by sampling location for *K* = 5. Individuals sampled within 80 km of each other are clustered together into the same pie. Pie size is scaled with the number of individuals. Individual ancestry proportions **B** before the hunting period, **C** during the hunting period, and **D** after the hunting period. C-Spitsbergen = Central Spitsbergen; S-Spitsbergen = South Spitsbergen; W-Spitsbergen = West Spitsbergen; E-Svalbard = East Svalbard.

In contrast, the spatio-temporal groups of ancient individuals are not strongly distinct from one another in the admixture analysis (Figure 2). All groups but pre-hunting Nordaustlandet (DS = 0) show ancestry from all five genetic clusters for *K*=5 (DS 0.593 - 0.890), irrespective of their geographic distance from one another. Diversity was maintained throughout the hunting period, apart from Wijdefjorden, which became less diverse (DS = 0.389). All ancient individuals share high proportions of ancestry with post-hunting Nordaustlandet (orange). This trend continues for higher values of *K* up until *K*=7. However, beginning with *K*=8, the similarity between post-hunting Nordaustlandet and ancient individuals (except for pre- hunting Nordaustlandet) largely disappears. Instead, we observed the emergence of a new cluster (brown) that is shared among all ancient, but none of the post-hunting individuals are assigned to it. Furthermore, all individuals from during-hunting Wijdefjorden are assigned exclusively to this cluster. As admixture analysis can be biased when having unequal sample sizes (Garcia-Erill & Albrechtsen, 2020; Puechmaille, 2016), we additionally performed an analysis with a reduced sample set with more equal sample sizes by reducing the number of contemporary samples. This analysis confirmed the results using the complete dataset (Fig. S16).

We found 30 distinct haplotypes from the mitogenome alignment of 108 individuals, and none of these haplotypes were shared between individuals in the pre- and post-hunting period (Figure 3). However, haplotype relatedness does not group per time period. Instead, post-hunting haplotypes were primarily at the periphery of the network, branching from different ancient ancestors located at the center of the network (Figure 3). Half of these 30 haplotypes belonged to 18 individuals from the ancient populations, while the other half belonged to the 90 individuals from the post-hunting populations. Each of the 12 individuals from the pre-hunting period had distinct haplotypes with up to 32 segregating sites. Individuals from the same region (i.e. East-Svalbard), had only 1-3 segregating sites despite the thousands of years separating the sampled individuals (e.g., East-Svalbard, 2037 years difference for only 1 segregation site difference). However, some individuals from the same region, living at approximately the same time (e.g., Central-Spitsbergen, 597 and 703 BP), have the most distant haplotypes for the given region. In the during-hunting period, four out of six individuals (66%) shared the same haplotype, and those were from the same region, Wijdefjorden. In the post-hunting period, we found up to 36 different segregation sites, forming seven distinct haplogroups separated by fewer than three mutations, where several individuals from the same region shared the same haplotype. However, within the same region, we also found very distant haplotypes with individuals more closely related to their common ancestral haplotype than to one another. For instance, in the Central-Spitsbergen region, modern haplotypes were either closely related to the haplotypes previously found in that same region, or previously found in Eastern-Svalbard. Therefore, haplotypes found today within a same region are more distant than haplotypes currently found across regions. Subsequently, although mitochondrial haplotype and nucleotide diversity were lower in the post-hunting than in the pre-hunting period, they have not been drastically reduced by hunting (Table 2).

**Table 2.** Measures of diversity statistics from the mitogenome analysis and genomic diversity. *N* = number of individuals; Num. haplo. = Number of haplotypes, Num. seg. sites. = Number of segregation sites, Hap. div = haplotype diversity and its standard deviation, Nuc. div = nuclear diversity, Med. het. = median heterozygosity, Nuc. div. = nucleotide diversity, *N_e_* = effective population size.

|  | Mitochondrial genome |  |  |  |  | Nuclear genome |  |  |  |
| --- | --- | --- | --- | --- | --- | --- | --- | --- | --- |
| Time period | *N* | Num. haplo. | Num. seg. sites | Hap. div. | Nuc. div. | *N* | Med. het. | Nuc. div. | *N<sub>e</sub>* |
| Pre-hunting | 12 | 12 | 32 | 1.00±5.79E-04 | 4.88E-04 | 5 | 0.00038 | 3.34E-04 | 5.21E+03 |
| During-hunting | 6 | 3 | 13 | 0.60±4.54E-02 | 3.03E-04 | 6 | 0.00031 | 3.35E-04 | 5.84E+03 |
| Ancient (Pre- and During-hunting) | 18 | 15 | 40 | 0.96±11.01E-04 | 4.84E-04 | 11 |  |  |  |
| Post-hunting | 90 | 15 | 36 | 0.85±2.42E-04 | 4.50E-04 | 90 | 0.00028 | 3.23E-04 | 4.66E+03 |
| All genomes | 108 | 30 | 56 | 0.90±1.80E-04 | 5.16E-04 | 101 |  |  |  |

**Figure 3.**
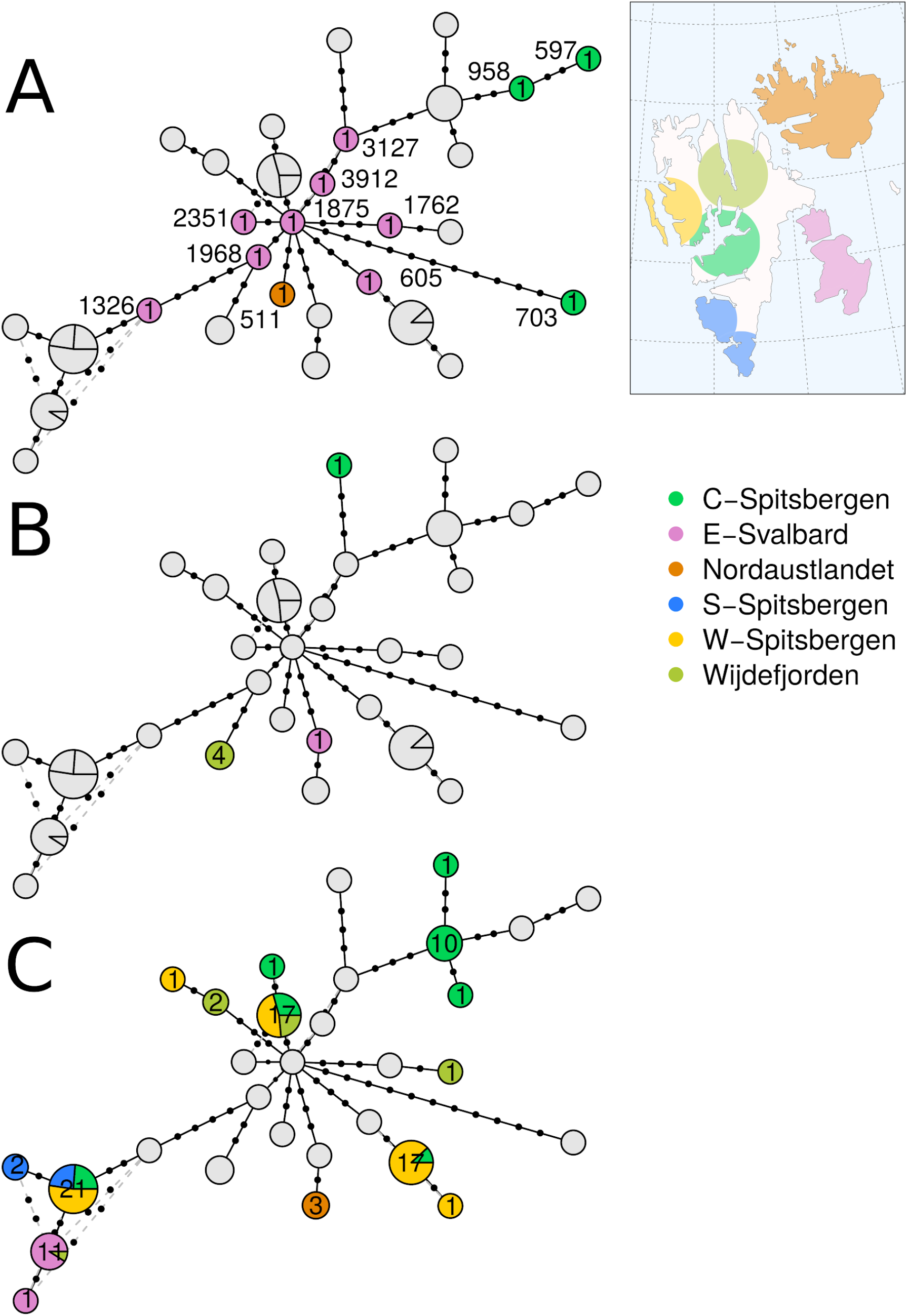
Haplotype network of Svalbard reindeer mitogenomes during three temporal hunting periods. **A: Pre-hunting. B: During-hunting. C: Post-hunting.** The number of individuals sharing the same haplotype is indicated in the center of the circle. Gray circles indicate the positioning of haplotypes that do not belong to the time period in focus. The diameter of the circles indicates the number of individuals sharing that haplotype and is scaled from 1 (i.e., min. haplotype number of 1) to 2 (i.e., max. haplotype number of 21). In the pre-hunting period, calibrated carbon-dated ages are reported as the median age probability in years Before Present (before 1950 AD). Calibrated age ranges are reported in Table 1. C-Spitsbergen = Central Spitsbergen; S-Spitsbergen = South Spitsbergen; W- Spitsbergen = West Spitsbergen; E-Svalbard = East Svalbard.

### 3.3 Temporal genomic differentiation

Heterozygosity decreased stepwise through the hunting periods, from the highest in the pre- hunting group to the lowest in the post-hunting group, although there was considerable overlap between the heterozygosity ranges of the three groups (see Figure 4). The median heterozygosities are 3.8 ✕ 10^-4^ for the pre-hunting period, 3.1 ✕ 10^-4^ for the during- hunting period, and 2.8 ✕ 10^-4^ for the post-hunting period. The differences in mean heterozygosity between during-hunting and the other two groups is not significant (pre- hunting: p=0.66, post-hunting: p=0.10, Mann-Whitney U test), while post-hunting heterozygosity is significantly lower than pre-hunting (p < 0.01, Mann-Whitney U test). The decrease in median heterozygosity from pre-hunting to during-hunting reindeer is 18.42%, and post-hunting reindeer have a 9.68% lower median heterozygosity than during-hunting reindeer. The median heterozygosity of post-hunting reindeer is 26.32% lower than that of pre-hunting reindeer. Mean effective population size *N_e_* was lower in the post-hunting population (4.66 ✕ 10^3^) than in the pre-hunting population (5.21 ✕ 10^3^), and highest in the during-hunting population (5.84 ✕ 10^3^). In the sliding window analysis, the mean nucleotide diversity across all windows was 3.34 ✕ 10^-4^ in the pre-hunting population, slightly higher at 3.35 ✕ 10^-4^ in the during-hunting population and reduced to 3.23 ✕ 10^-4^ n the post-hunting population.

**Figure 4.**
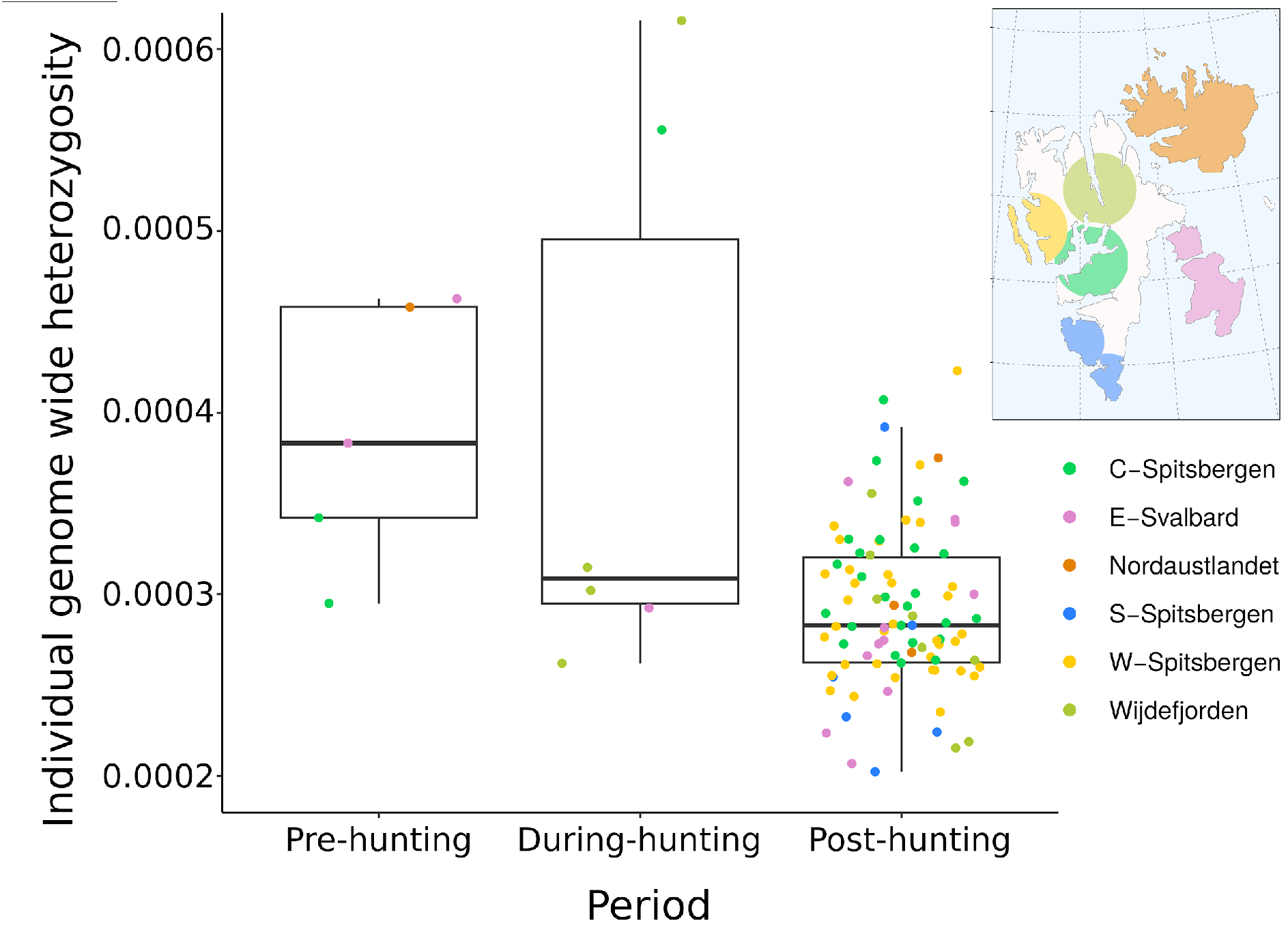
Individual genome-wide heterozygosity by time period. The points are horizontally scattered to increase visibility. Thick horizontal line in the boxplot represents the median, the lower and upper box bound represent the 25 and 75 percentile, respectively. C- Spitsbergen = Central Spitsbergen; S-Spitsbergen = South Spitsbergen; W-Spitsbergen = West Spitsbergen; E-Svalbard = East Svalbard.

There were a total of 217,247 windows identified by the sliding window analysis, of which 183 (0.1%) were *F_ST_* outlier windows as defined by a Z-score ≥ 6 (Figure 5). The mean value of weighted pairwise *F_ST_* was 1.91 ✕ 10^-2^ for the pre-hunting and during-hunting populations, 3.27 ✕ 10^-2^ for the pre-hunting and post-hunting populations, and 5.06 ✕ 10^-2^ for the during-hunting and post-hunting populations. The mean number of sites across all windows was 8,649, and the mean number of sites within outlier windows was 8,843. To assess whether windows were under positive selection, we performed neutrality tests to compare Fay & Wu’s *H* between during-hunting outlier and non-outlier windows, as well as between post-hunting outlier and non-outlier windows. Comparison shows that *H* is much lower and more broadly distributed in ancient outlier windows compared to all the other groups (Figure 5 and Figure S18).

**Figure 5:**
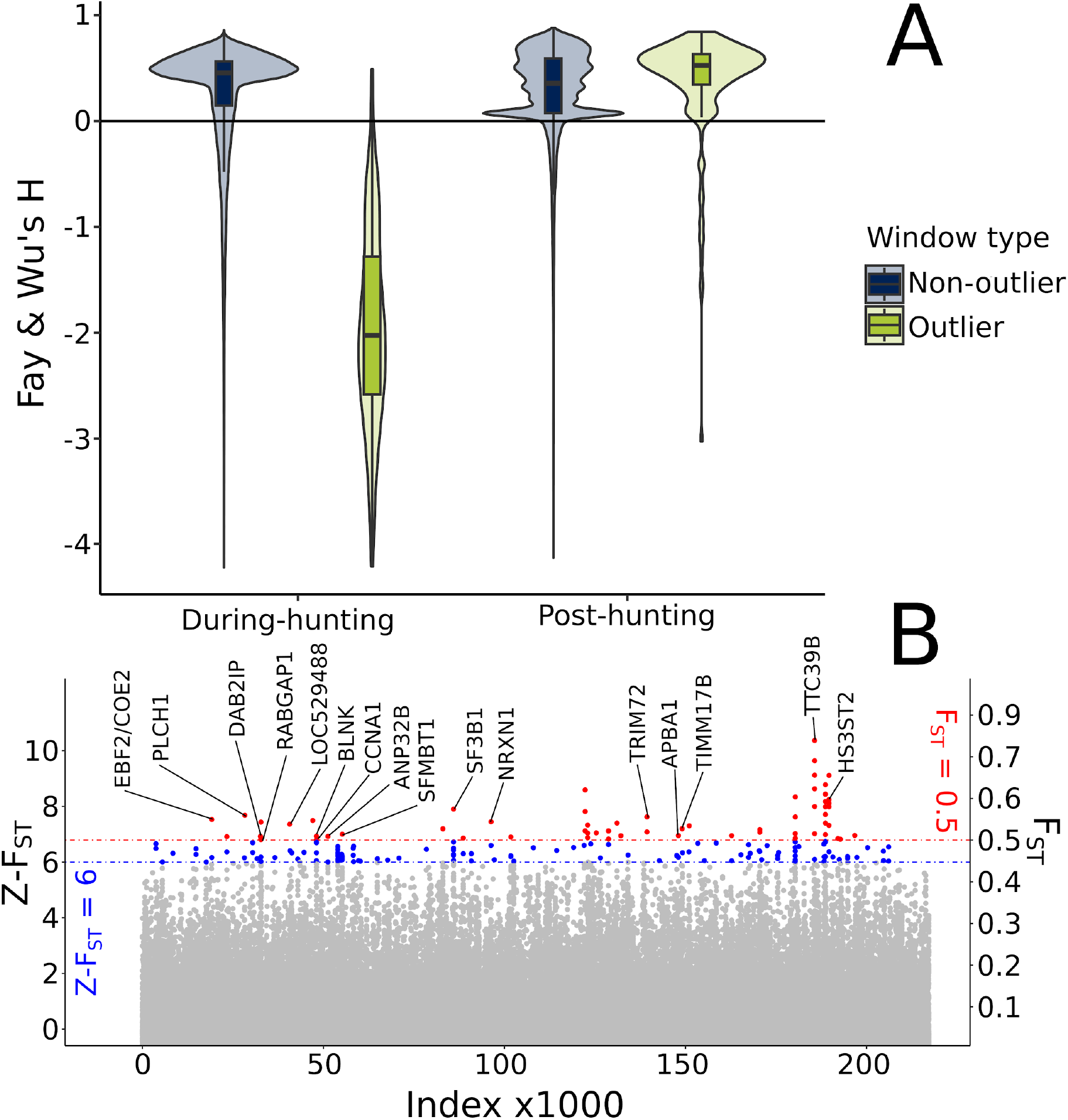
Outlier genomic windows and their genes. **A:** Comparison of Fay & Wu’s H in genomic windows as measure of neutrality. Here, *H* was measured in genomic regions (windows) in genomes of individuals living during- (left) and post-hunting (right). Regions that strongly diverged (z-F_ST_ ≥ 6) between the during-hunting and post-hunting time periods are called ‘outliers’ (green), those that are not ‘non-outliers’ (blue). Each violin shows the distribution of *H* in each type/period pair respectively. **B:** Manhattan plot of genomic windows, highlighting windows that cross the threshold of Z-F_ST_ greater than or equal to 6 (blue) and F_ST_ of greater than or equal to 0.5 (red). High F_ST_ windows that intersect with described genes are labeled with the gene name.

To explore any functional genetic changes that may have resulted from overhunting and near-extirpation of Svalbard reindeer, we further investigated those outlier genomic regions with extraordinarily high *F_ST_* (*F_ST_* ≥ 0.5) as measured between the during-hunting and post-hunting populations, as well as the identity and functions of the genes therein (Table S4). We identified 50 high-divergence outlier windows which intersect with genes, of which 34 are unique annotated genes (Taylor et al., 2019). A blastp search against the *Bos taurus* proteome (see Materials and Methods) matched 30 of these 34 genes to 29 unique *Bos taurus* proteins (Table S4). Of these 29, 17 are predicted proteins, nine are further inferred from homology, and four have evidence at transcriptome level. For 17 of these proteins, the associated coding gene is known in the UniProt database. The known coding genes are *PLCH1*, *TRIM72* (*MG53*), *TIMM17B*, *ANP32B*, *APBA1*, *BLNK*, *CCNA1, DAB2IP*, *EBF2/COE2*, *HS3ST2*, *LOC529488*, *NRXN1*, *RABGAP1*, *SF3B1*, *SFMBT1*, *STRBP*, and *TTC39B*.

## 4. Discussion

Here, by combining modern and the first ancient genomes (up to 4000 BP), we have described and compared the population structure and genetic changes of Svalbard reindeer before, during and after an intense period of anthropogenic harvest . We have shown that the population collapse due to overharvest decreased nuclear genomic diversity as well as effective population size (Figure 4, Table 2). The overhunting event also resulted in a shift of genetic diversity across the entire archipelago (Figure 1 and 2), suggesting weaker population substructure before human presence and harvest. Mitogenome analysis also revealed a significant loss, with none of the pre-hunting haplotypes occurring today (Figure 3, Table 2). Hence, post-hunting haplotypes formed their own haplogroups, which were more closely linked to ancient haplotypes from different regions of Svalbard than to the ancient haplotypes from their respective sampling locations. Gene selection analyses indicate pronounced genetic drift during and post-hunting periods rather than natural selection (Figure 5).

### 4.1 Major changes in population structure and differentiation

Our analysis of population structure based on whole nuclear genomes revealed substantial differences between historical/ancient samples and present-day Svalbard reindeer (Burnett et al., 2022; Peeters et al., 2020). We identified *K* = 5 as the optimal value of *K* with the delta-*K* method, but in light of potential sampling bias and strong population structure both in space and time, we elect to discuss higher values of *K* as well. In contrast to the high degree of nuclear genetic differentiation in the post-hunting populations, which show a near-perfect correspondence of geographical and genetic grouping, Svalbard reindeer did not form distinct genetic clusters based on location prior to and during hunting. Instead, all genomes but one (from the most distant individual from the isolated island of Nordaustlandet, *n* = 1) had higher nuclear genetic diversity in the pre-hunting period. These indicate a scenario where ancient Svalbard reindeer formed a single genetically diverse, continuous and panmictic population in the past, a similar situation as for the Iberian lynx (*Lynx pardinus* (Temminck, 1827)), another species with severe history of overharvest (Casas-Marce et al., 2017). Pre-hunting Svalbard reindeer individuals showed affinity to all post-hunting metapopulations for *K*=2 to *K*=7 which suggests high levels of gene-flow between different geographic regions in ancient Svalbard or even a single panmictic population . However, the level of gene-flow was likely not uniform across Svalbard. Individuals from modern Nordaustlandet were most similar to their ancient counterpart for *K*<8, the reason for which could be that the size of this remnant population remained low, while other remnant populations underwent rapid population growth and geographic expansion (Le Moullec et al. 2019). This interpretation is supported by the PCA, which places modern Nordaustlandet closer than other modern samples to the ancient samples. Ancient Wijdefjorden has high affinity to ancient and modern Nordaustlandet for *K*<8, however, at *K*=8 they lose all affinity with modern samples and become assigned to an ancient-only cluster, with high, but reduced affinity to ancient Nordaustlandet. The other ancient samples are partly assigned to this ancient-only cluster, but retain relatively high ancestry to genetic clusters also found in modern samples at higher values of *K*. These populations differentiated from each other through time. This change happened between the during-hunting and post-hunting groups rather than between the pre-hunting and during-hunting groups, which cover a much longer time-span, suggesting that the change was related to overharvest.

We observed a gradual decrease in Svalbard reindeer genome heterozygosity over the three time periods. Median heterozygosity was reduced by 26% over the time spanning the pre- hunting and the post-hunting periods. This decrease is congruent with results obtained from previous studies comparing modern to historic/ancient vertebrates that underwent large- scale population declines due to anthropogenic near-extirpation events, for example the white rhinoceros (*Ceratotherium simum* (Burchell, 1817))((Sánchez-Barreiro et al., 2021)), alpine ibex (*Capra ibex*, Linnaeus, 1758) ((Robin et al., 2022)), eastern gorilla (*Gorilla beringe*i Matschie, 1903)(van der Valk et al., 2019) and iberian lynx (Casas-Marce et al., 2017). The Svalbard reindeer’s decrease of heterozygosity over time is of similar severity to that measured following drastic population declines in the Iberian lynx (heterozygosity reduction of 10% based on microsatellite data, (Casas-Marce et al., 2017) and in two over- harvested populations of white rhinoceros (heterozygosity reductions of 10% and 37% respectively, (Sánchez-Barreiro et al., 2021). A study by (van der Valk et al., 2019) reported a 20% decrease of heterozygosity in Grauer’s gorillas but only a 3% decrease in mountain gorillas. It is important to note that prior to overharvesting, the Svalbard reindeer population genome-wide heterozygosity was already low, perhaps because of a strong bottleneck when colonizing Svalbard.

Our measurements of temporal change in mitogenome diversity did not suggest a strong decrease in genetic diversity. The relatively high diversity of modern haplogroups (*n* = 7) contrasts with other ungulate species heavily hunted in the past, like the Alpine ibex, where only two modern haplogroups are now widespread across their range (Robin et al., 2022).

Contrary to the Alpine ibex, in which mitogenome nucleotide diversity was reduced by ∼79 % (6.38 ✕ 10^-4^) from the pre-hunting to the post-hunting period, Svalbard reindeer experienced a reduction of only ∼8% (0.38 ✕ 10^-4^). However, this difference could be explained by the fact that after near-extirpation, the Svalbard reindeer survived in four remnant populations, as opposed to the Alpine ibex, which survived in only a single remnant population. During the intermediate hunting-period, the mitogenome nucleotide diversity surprisingly dropped, yet this is likely due to the low sample size. In this period, large individual variation existed, where two of the six individuals had the highest genome-wide heterozygosity observed in our dataset, while the other four were among the lowest values. The uncertainty related to the carbon-dating calibration scale prior to the industrial era, which coincides with our during-intermediate hunting period (1500-1950 CE), restricts the temporal resolution of this period. However, previous reports documented a harvest peak in the early 1900s CE (Hoel, 1916; Lønø, 1959), just prior to the legal protection of Svalbard reindeer in 1925. Thus, individuals with high genetic diversity are likely to have lived prior to this harvesting peak.

### Potential climate change impacts during recovery

Accounting for the fact that current summer temperature has already increased by 1.5-2°C since the reference period of 1912-2012 CE (Isaksen et al., 2022; van der Bilt et al., 2019), the older reindeer specimens from our collection (ages of 3000-4000 BP) experienced summer temperatures that were similar to the present-day climate. From this period and until harvesting started ∼1500 CE, the relatively stable climate became gradually cooler with sea-ice cover persisting year-round (Werner et al., 2016), likely acting as dispersal corridors for reindeer. These conditions likely favored a higher degree of admixture between populations.

The current population structuring is a consequence of overharvesting and recovery occurring during a period of pronounced climate warming. Our results are congruent with earlier findings based on microsatellite data and nuclear whole genome sequencing where present-day population structuring reflects the recolonization patterns originating from the four locations that escaped extirpation (Burnett et al., 2022; Peeters et al., 2020). In addition to the sedentary behavior of the Svalbard reindeer, the effect of natural barriers for dispersal and gene flow has increased during the recovery period, as sea-ice cover decreased (Peeters et al., 2020). Despite this, Svalbard reindeer were capable to disperse naturally and through reintroduction events to all suitable habitats on Svalbard within a century after protection (Le Moullec et al., 2019). The increased isolation due to sea-ice cover decline may have been partly counteracted by recent increases in frequency of rain-on-snow events resulting in winters with poor feeding conditions (Peeters et al., 2019). That is, such extreme events can sometimes aid recolonization by ‘pushing’ reindeer to disperse to neighboring islands or peninsulas where they were previously extirpated (Hansen et al., 2011). This may have accelerated the natural recolonization process, likely through a stepping-stone process that conserve geographic genetic structuring, but also likely further decreased genetic diversity in the peripheral recolonized populations. Still, the rapid increase in temperatures in Svalbard (Isaksen et al., 2022) and associated sea-ice decline likely restrict dispersal more nowadays than over the millennia prior to anthropogenic disturbance.

### Stochastic changes within diverse gene families

For the pre-hunting and during-hunting populations, we observed strongly negative values of Fay & Wu’s *H* within *F_ST_* outlier windows, whereas these *F_ST_* outlier windows had generally positive *H* values in the post-hunting population . This indicates that these genomic regions were experiencing positive selection in the pre-hunting and during-hunting periods, but now evolve mainly under genetic drift in the extant post-hunting population, rather than by natural selection (Fay & Wu, 2000). Conceivably, selection pressures from the extreme Arctic environment were the major force acting on Svalbard reindeer under natural conditions prior to anthropogenic disturbance. However, genetic drift can rapidly and stochastically alter allele frequencies, including at loci that were previously conserved under strong selective pressures, especially in small populations (Bortoluzzi et al., 2020). Future research would be needed to confirm whether there are functional consequences, e.g. affecting fecundity, survival, or behavior.

In that context it is relevant to explore which genes were most affected by genetic drift following overhunting, since they may have functional relevance for the health and conservation of the present-day Svalbard reindeer population. Owing to the stochastic nature of genetic drift, the affected genes and their coded proteins are involved in a great variety of biological functions. However, among the candidate genes we identified are some that play key roles in circadian rhythm regulation, fat storage, the immune system, the nervous system, and basic neurological functions. One of those genes in a region highly affected by genetic drift is an ortholog of *PLCH1* (phospholipase-C eta1) that encodes the protein *Phosphoinositide phospholipase C (PLC). PLC* is involved in cAMP-responsive element-binding protein (*CREB*) mediated gene transcription, which activates the transcription of the genes *PER1* and *PER2* (among others) in response to light (Colwell, 2011). *PER2* has reindeer-specific mutations and has been linked to the lack of circadian rhythmicity in reindeer (Lin et al., 2019). The loss of a day/night controlled internal biological clock is considered an important adaptation of reindeer to high Arctic environments, where daylight conditions do not change for extended periods of the year (Lin et al., 2019; van Oort et al., 2005).

We also identified genes that relate to lipid metabolism. *TRIM72* / *MG53* is a multifunctional gene primarily involved in cell membrane repair and tissue regeneration, but it also has been linked to insulin resistance and related metabolic abnormalities, including obesity (Z. Li et al., 2021; Song et al., 2013; Y. Zhang et al., 2017). *APBA1* is involved in insulin secretion (K. Zhang et al., 2021). *TIMM17B* is a protein that mediates inner mitochondrial membrane transport, and it may be related to insulin resistance and obesity in human populations (Dubé et al., 2020). *TTC39B* has been found to be involved in lipid metabolism and coronary artery diseases (Teslovich et al., 2010). *EBF2/COE2* encodes *Early B-Cell factor 2 (Ebf2)* which is highly involved in the formation of brown adipose tissue (Wang et al., 2014). The Svalbard reindeer’s overwinter survival is dependent on its ability to build up high ratios of body fat relative to total body mass over the very short snow-free season (Trondrud et al., 2021), normally lasting for about three to four months. It is not unlikely that genes linked to excessive fat accumulation (i.e. obesity) in other species are the underlying genes that support rapid fattening in Svalbard reindeer. The outlier window containing *EBF2/COE2* was the sole window with a negative *H* value in the post-hunting population, suggesting it may be under strong purifying selection. This could indicate that this gene is of importance for survival and therefore influenced by comparatively stronger purifying selection and weaker genetic drift than other genes.

Furthermore, some of these genes we found are involved in spermatogenesis and therefore might play a role in Svalbard reindeer fertility. *CCNA1* encodes the protein *Cyclin A1*, which plays a vital role in male mammalian meiosis and spermatogenesis, and loss of *CCNA1* causes infertility in males, as demonstrated in mice (Liu et al., 1998). *SFBT1* and *STRBP* are genes involved in spermatogenesis as well (Pires-daSilva et al., 2001; J. Zhang et al., 2013). We also identified *Testis-specific Y-encoded-like protein 6*, which in humans is encoded by the gene *TSPYL6*, and is expressed only in the testes and involved in spermatogenesis as well (Uhlén et al., 2015). *BLNK* is the coding gene for the B-cell receptor that is related to B-cell function and development and therefore likely plays a role in the Svalbard reindeer immune system (Fu et al., 1998).

The genes we found that are related to neuronal development are *NRXN1* and *LOC529488. LOC529488* encodes *Glutamate decarboxylase 1* (*GAD1*), which is responsible for production of the inhibitory neurotransmitter GABA (Fenalti et al., 2007). *Neurexin-1 (NRXN1)* from the Neurexin protein family of synaptic adhesion molecules is involved in GABA release, and mutations in it have been linked to developmental and neuropsychiatric disorders (Hu et al., 2019; Missler et al., 2003).

### 4.2 Implications for conservation

Our study supplements a growing body of research utilizing temporal datasets to assess genomic health of threatened and endemic species. Our analyses indicate that historical overharvest has, in addition to the previously reported population size reduction (Lønø, 1959), decreased overall genomic diversity in Svalbard reindeer. While subsequent protection of the Svalbard reindeer and its habitat has rapidly (i.e., rapidly in an evolutionary context) facilitated the recolonization of the archipelago (Le Moullec et al., 2019), the current level of genome erosion could make this subspecies particularly vulnerable to future climate and environmental changes and associated demographic stochasticity, especially if inbreeding levels remain high (Burnett et al., 2022; Peeters et al., 2020). Our findings support the view that census data on population abundances alone is not robust enough to assess the conservation status of populations recovering from overharvest or other anthropogenic stressors. Genomic monitoring, especially when incorporating a temporal component derived from ancient DNA, can help capture a more complete understanding (Jensen et al., 2022).

In combination with such genomic monitoring, translocations have been suggested as an effective conservation measure (Bertola et al., 2022; Bubac et al., 2019). In contrast to other species (e.g., Iberian and Alpine ibex; (Grossen et al., 2018, 2020)), translocation events have strongly contributed to limit the genetic diversity loss in Svalbard reindeer caused by overharvesting, likely because the translocated individuals came from the population with the highest genetic diversity levels (Burnett et al., 2022). Nevertheless, a population’s adaptive potential relies not only on their overall level of genetic diversity, but also on functional diversity. When the genetic diversity of a species is low and there has been significant genetic turnover due to genetic drift rather than natural selection 一as in Svalbard reindeer 一the species’ capacity to evolve with climate change has possibly been reduced.

## Acknowledgements

This study was primarily financed by the Research Council of Norway FRIMEDBIO awards 325589 to MDM and 276080 to BBH, the Svalbard Environmental Protection Fund awards 14/137 and 15/105 to BBH, and internal NTNU funding to MDM. We also thank M. Thomas P. Gilbert for his valuable financial support of the computational analyses. We are grateful to Marie-Josée Nadeau and Martin Seiler at the National Laboratory of Age Determination, NTNU and to Love Dalén for support and advice early in the project period. We acknowledge the following sequencing centers and associates for their invaluable services: Novogene (Cambridge, UK), the Norwegian National Sequencing Centre (Oslo, Norway), and the NTNU Genomics Core Facility (Trondheim, Norway). Some computational analyses were performed on resources provided by the Danish National Life Science Supercomputing Center (Computerome) and the National Infrastructure for High Performance Computing and Data Storage in Norway (UNINETT Sigma2) under projects NN8052K and NS8052K.

## Data Accessibility Statement

Raw sequence data generated for this study has been archived at the European Nucleotide Archive under accession code PRJEB60484. Sample metadata can be accessed in the supplementary material (Table S3).

## Author Contributions

FLK, MLM wrote the paper with input from all authors

FLK, MLM analyzed data with input from MDM, VCB, JCB, HB

FLK, MLM, MDM, VCB, ND, BP contributed to interpreting the results

MRE, VCB, MDM, BP did ancient DNA lab work

MLM, BBH, JR performed ancient sample acquisition and preparation BBH, MLM, MDM had the original idea for the study

FLK, MLM, MDM, BBH, JR, BP, VCB designed the study MDM, VCB supervised the paleogenomic work

MDM, BBH funded the study

## References

Aanes, R., Saether, B.-E., & Øritsland, N. A. (2000). Fluctuations of an introduced population of Svalbard reindeer: the effects of density dependence and climatic variation. Ecography, 23(4), 437–443. 10.1111/j.1600-0587.2000.tb00300.x

Albon, S. D., Irvine, R. J., Halvorsen, O., Langvatn, R., Loe, L. E., Ropstad, E., Veiberg, V., van der Wal, R., Bjørkvoll, E. M., Duff, E. I., Hansen, B. B., Lee, A. M., Tveraa, T., & Stien, A. (2017). Contrasting effects of summer and winter warming on body mass explain population dynamics in a food-limited Arctic herbivore. Global Change Biology, 23(4), 1374–1389. 10.1111/gcb.13435

Allendorf, F. W., & Hard, J. J. (2009). Human-induced evolution caused by unnatural selection through harvest of wild animals. Proceedings of the National Academy of Sciences of the United States of America, 106 Suppl 1(Suppl 1), 9987–9994. 10.1073/pnas.0901069106

Altschul, S. F., Gish, W., Miller, W., Myers, E. W., & Lipman, D. J. (1990). Basic local alignment search tool. Journal of Molecular Biology, 215(3), 403–410. 10.1016/S0022-2836(05)80360-2

Bana, N. Á., Nyiri, A., Nagy, J., Frank, K., Nagy, T., Stéger, V., Schiller, M., Lakatos, P., Sugár, L., Horn, P., Barta, E., & Orosz, L. (2018). The red deer Cervus elaphus genome CerEla1.0: sequencing, annotating, genes, and chromosomes. Molecular Genetics and Genomics: MGG, 293(3), 665–684. 10.1007/s00438-017-1412-3

Bertola, L. D., Miller, S. M., Williams, V. L., Naude, V. N., Coals, P., Dures, S. G., Henschel, P., Chege, M., Sogbohossou, E. A., Ndiaye, A., Kiki, M., Gaylard, A., Ikanda, D. K., Becker, M. S., & Lindsey, P. (2022). Genetic guidelines for translocations: Maintaining intraspecific diversity in the lion (Panthera leo). Evolutionary Applications, 15(1), 22–39. 10.1111/eva.13318

Bortoluzzi, C., Bosse, M., Derks, M. F. L., Crooijmans, R. P. M. A., Groenen, M. A. M., & Megens, H.-J. (2020). The type of bottleneck matters: Insights into the deleterious variation landscape of small managed populations. Evolutionary Applications, 13(2), 330–341. 10.1111/eva.12872

Bowyer, R. T., Boyce, M. S., & Goheen, J. R. (2019). Conservation of the world’s mammals: status, protected areas, community efforts, and hunting. Journal of. https://academic.oup.com/jmammal/article-abstract/100/3/923/5498008

Bubac, C. M., Johnson, A. C., Fox, J. A., & Cullingham, C. I. (2019). Conservation translocations and post-release monitoring: Identifying trends in failures, biases, and challenges from around the world. Biological Conservation, 238, 108239. 10.1016/j.biocon.2019.108239

Burnett, H. A., Bieker, V. C., Le Moullec, M., Peeters, B., Rosvold, J., Pedersen, Å. Ø., Dalén, L., Loe, L. E., Jensen, H., Hansen, B. B., & Martin, M. D. (2022). Contrasting genomic consequences of anthropogenic reintroduction and natural recolonisation in high-arctic wild reindeer. In bioRxiv (p. 2022.11.25.517957). 10.1101/2022.11.25.517957

Byun, S. A., Koop, B. F., & Reimchen, T. E. (2002). Evolution of the Dawson caribou (Rangifer tarandus dawsoni). Canadian Journal of Zoology, 80(5), 956–960. 10.1139/z02-062

CAFF. (2013). Arctic Biodiversity Assessment: Status and trends in Arctic biodiversity. Conservation of Arctic Flora and Fauna (H. Meltofte (ed.)). CAFF International Secretariat. https://play.google.com/store/books/details?id=-P_FngEACAAJ

Carøe, C., Gopalakrishnan, S., Vinner, L., Mak, S. S. T., Sinding, M. H. S., Samaniego, J. A., Wales, N., Sicheritz-Pontén, T., & Gilbert, M. T. P. (2018). Single-tube library preparation for degraded DNA. Methods in Ecology and Evolution / British Ecological Society, 9(2), 410–419. 10.1111/2041-210X.12871

Casas-Marce, M., Marmesat, E., Soriano, L., Martínez-Cruz, B., Lucena-Perez, M., Nocete, F., Rodríguez-Hidalgo, A., Canals, A., Nadal, J., Detry, C., Bernáldez-Sánchez, E., Fernández-Rodríguez, C., Pérez-Ripoll, M., Stiller, M., Hofreiter, M., Rodríguez, A., Revilla, E., Delibes, M., & Godoy, J. A. (2017). Spatiotemporal Dynamics of Genetic Variation in the Iberian Lynx along Its Path to Extinction Reconstructed with Ancient DNA. Molecular Biology and Evolution, 34(11), 2893–2907. 10.1093/molbev/msx222

Chang, C. C., Chow, C. C., Tellier, L. C., Vattikuti, S., Purcell, S. M., & Lee, J. J. (2015). Second-generation PLINK: rising to the challenge of larger and richer datasets. GigaScience, 4, 7. 10.1186/s13742-015-0047-8

Collard, R.-C., Dempsey, J., & Holmberg, M. (2020). Extirpation despite regulation? Environmental assessment and caribou. Conservation Science and Practice, 2(4). 10.1111/csp2.166

Colwell, C. S. (2011). Linking neural activity and molecular oscillations in the SCN. Nature Reviews. Neuroscience, 12(10), 553–569. 10.1038/nrn3086

Crema, E. R., & Bevan, A. (2021). INFERENCE FROM LARGE SETS OF RADIOCARBON DATES: SOFTWARE AND METHODS. Radiocarbon, 63(1), 23–39. 10.1017/RDC.2020.95

Derocher, A. E., Wiig, Ø., & Bangjord, G. (2000). Predation of Svalbard reindeer by polar bears. Polar Biology, 23(10), 675–678. 10.1007/s003000000138

Díez-Del-Molino, D., Sánchez-Barreiro, F., Barnes, I., Gilbert, M. T. P., & Dalén, L. (2018). Quantifying Temporal Genomic Erosion in Endangered Species. Trends in Ecology & Evolution, 33(3), 176–185. 10.1016/j.tree.2017.12.002

Drummond, A. J., Rambaut, A., Shapiro, B., & Pybus, O. G. (2005). Bayesian coalescent inference of past population dynamics from molecular sequences. Molecular Biology and Evolution, 22(5), 1185–1192. 10.1093/molbev/msi103

Dubé, J. J., Collyer, M. L., Trant, S., Toledo, F. G. S., Goodpaster, B. H., Kershaw, E. E., & DeLany, J. P. (2020). Decreased Mitochondrial Dynamics Is Associated with Insulin Resistance, Metabolic Rate, and Fitness in African Americans. The Journal of Clinical Endocrinology and Metabolism, 105(4), 1210–1220. 10.1210/clinem/dgz272

Dussex, N., Alberti, F., Heino, M. T., Olsen, R.-A., van der Valk, T., Ryman, N., Laikre, L., Ahlgren, H., Askeyev, I. V., Askeyev, O. V., Shaymuratova, D. N., Askeyev, A. O., Döppes, D., Friedrich, R., Lindauer, S., Rosendahl, W., Aspi, J., Hofreiter, M., Lidén, K., … Díez-Del-Molino, D. (2020). Moose genomes reveal past glacial demography and the origin of modern lineages. BMC Genomics, 21(1), 854. 10.1186/s12864-020-07208-3

Evanno, G., Regnaut, S., & Goudet, J. (2005). Detecting the number of clusters of individuals using the software STRUCTURE: a simulation study. Molecular Ecology, 14(8), 2611–2620. 10.1111/j.1365-294X.2005.02553.x

Fay, J. C., & Wu, C. I. (2000). Hitchhiking under positive Darwinian selection. Genetics, 155(3), 1405–1413. 10.1093/genetics/155.3.1405

Fenalti, G., Law, R. H. P., Buckle, A. M., Langendorf, C., Tuck, K., Rosado, C. J., Faux, N. G., Mahmood, K., Hampe, C. S., Banga, J. P., Wilce, M., Schmidberger, J., Rossjohn, J., El-Kabbani, O., Pike, R. N., Smith, A. I., Mackay, I. R., Rowley, M. J., & Whisstock, J. C. (2007). GABA production by glutamic acid decarboxylase is regulated by a dynamic catalytic loop. Nature Structural & Molecular Biology, 14(4), 280–286. 10.1038/nsmb1228

Festa-Bianchet, M., Ray, J. C., Boutin, S., Côté, S. D., & Gunn, A. (2011). Conservation of caribou (Rangifer tarandus) in Canada: an uncertain future. Canadian Journal of Zoology, 89(5), 419–434. 10.1139/z11-025

Flagstad, Ø., Kvalnes, T., Røed, K. H., Våge, J., Strand, O., & Sæther, B.-E. (2022). Genetisk levedyktig villreinbestand på Hardangervidda (No. 2176). Norsk institutt for naturforskning. https://brage.nina.no/nina-xmlui/handle/11250/3020843

Frankham, R. (2005). Genetics and extinction. Biological Conservation, 126(2), 131–140. 10.1016/j.biocon.2005.05.002

Frankham, R., Ballou, S. E. J., Briscoe, D. A., & Ballou, J. D. (2002). Introduction to Conservation Genetics. Cambridge University Press. https://play.google.com/store/books/details?id=F-XB8hqZ4s8C

Fu, C., Turck, C. W., Kurosaki, T., & Chan, A. C. (1998). BLNK: a central linker protein in B cell activation. Immunity, 9(1), 93–103. 10.1016/s1074-7613(00)80591-9

Garcia-Erill, G., & Albrechtsen, A. (2020). Evaluation of model fit of inferred admixture proportions. Molecular Ecology Resources, 20(4), 936–949. 10.1111/1755-0998.13171

Gravlund, P., Meldgaard, M., Pääbo, S., & Arctander, P. (1998). Polyphyletic origin of the small-bodied, high-arctic subspecies of tundra reindeer (Rangifer tarandus). Molecular Phylogenetics and Evolution, 10(2), 151–159. 10.1006/mpev.1998.0525

Grossen, C., Biebach, I., Angelone-Alasaad, S., Keller, L. F., & Croll, D. (2018). Population genomics analyses of European ibex species show lower diversity and higher inbreeding in reintroduced populations. Evolutionary Applications, 11(2), 123–139. 10.1111/eva.12490

Grossen, C., Guillaume, F., Keller, L. F., & Croll, D. (2020). Purging of highly deleterious mutations through severe bottlenecks in Alpine ibex. Nature Communications, 11(1), 1001. 10.1038/s41467-020-14803-1

Hansen, B. B., Aanes, R., Herfindal, I., Kohler, J., & Saether, B.-E. (2011). Climate, icing, and wild arctic reindeer: past relationships and future prospects. Ecology, 92(10), 1917– 1923. 10.1890/11-0095.1

Hansen, B. B., Pedersen, Å. Ø., Peeters, B., Le Moullec, M., Albon, S. D., Herfindal, I., Sæther, B., Grøtan, V., & Aanes, R. (2019). Spatial heterogeneity in climate change effects decouples the long-term dynamics of wild reindeer populations in the high Arctic. Global Change Biology, March, 3656–3668. 10.1111/gcb.14761

Harismendy, O., Kim, J., Xu, X., & Ohno-Machado, L. (2019). Evaluating and sharing global genetic ancestry in biomedical datasets. Journal of the American Medical Informatics Association: JAMIA, 26(5), 457–461. 10.1093/jamia/ocy194

Heckeberg, N., & Wörheide, G. (2019). A comprehensive approach towards the systematics of Cervidae. 10.7287/peerj.preprints.27618

Hoel, A. (1916). Hvorfra er Spitsbergrenen kommet? Naturen.

Hu, Z., Xiao, X., Zhang, Z., & Li, M. (2019). Genetic insights and neurobiological implications from NRXN1 in neuropsychiatric disorders. Molecular Psychiatry, 24(10), 1400–1414. 10.1038/s41380-019-0438-9

Isaksen, K., Nordli, Ø., Ivanov, B., Køltzow, M. A. Ø., Aaboe, S., Gjelten, H. M., Mezghani, A., Eastwood, S., Førland, E., Benestad, R. E., Hanssen-Bauer, I., Brækkan, R., Sviashchennikov, P., Demin, V., Revina, A., & Karandasheva, T. (2022). Exceptional warming over the Barents area. Scientific Reports, 12(1), 9371. 10.1038/s41598-022-13568-5

IUCN. (2020). The IUCN red list of threatened species. Version 2020-1. IUCN Red List of Threatened Species (2020).

Jensen, E. L., Díez-Del-Molino, D., Gilbert, M. T. P., Bertola, L. D., Borges, F., Cubric-Curik, V., de Navascués, M., Frandsen, P., Heuertz, M., Hvilsom, C., Jiménez-Mena, B., Miettinen, A., Moest, M., Pečnerová, P., Barnes, I., & Vernesi, C. (2022). Ancient and historical DNA in conservation policy. Trends in Ecology & Evolution, 37(5), 420–429. 10.1016/j.tree.2021.12.010

Johansen, B. E., Karlsen, S. R., & Tømmervik, H. (2012). Vegetation mapping of Svalbard utilising Landsat TM/ETM+ data. The Polar Record, 48(1), 47–63. 10.1017/s0032247411000647

Jónsson, H., Ginolhac, A., Schubert, M., Johnson, P. L. F., & Orlando, L. (2013). mapDamage2.0: fast approximate Bayesian estimates of ancient DNA damage parameters. Bioinformatics, 29(13), 1682–1684. 10.1093/bioinformatics/btt193

Kircher, M., Sawyer, S., & Meyer, M. (2012). Double indexing overcomes inaccuracies in multiplex sequencing on the Illumina platform. Nucleic Acids Research, 40(1), e3. 10.1093/nar/gkr771

Kohn, M. H., Murphy, W. J., Ostrander, E. A., & Wayne, R. K. (2006). Genomics and conservation genetics. Trends in Ecology & Evolution, 21(11), 629–637. 10.1016/j.tree.2006.08.001

Korneliussen, T. S., Albrechtsen, A., & Nielsen, R. (2014). ANGSD: Analysis of Next Generation Sequencing Data. BMC Bioinformatics, 15, 356. 10.1186/s12859-014-0356-4

Kumar, S., & Subramanian, S. (2002). Mutation rates in mammalian genomes. Proceedings of the National Academy of Sciences of the United States of America, 99(2), 803–808. 10.1073/pnas.022629899

Kvie, K. S., Heggenes, J., Anderson, D. G., Kholodova, M. V., Sipko, T., Mizin, I., & Røed, K. H. (2016). Colonizing the high arctic: Mitochondrial DNA reveals common origin of Eurasian archipelagic reindeer (Rangifer tarandus). PloS One, 11(11), 1–15. 10.1371/journal.pone.0165237

Laikre, L., Allendorf, F. W., Aroner, L. C., Baker, C. S., Gregovich, D. P., Hansen, M. M., Jackson, J. A., Kendall, K. C., McKelvey, K., Neel, M. C., Olivieri, I., Ryman, N., Schwartz, M. K., Bull, R. S., Stetz, J. B., Tallmon, D. A., Taylor, B. L., Vojta, C. D., Waller, D. M., & Waples, R. S. (2010). Neglect of Genetic Diversity in Implementation of the Convention of Biological Diversity. Conservation Biology: The Journal of the Society for Conservation Biology, 24(1), 86–88. http://www.jstor.org/stable/40419633

Lande, R., Engen, S., & Sæther, B.-E. (2003). Stochastic Population Dynamics in Ecology and Conservation. Oxford University Press. https://play.google.com/store/books/details?id=6KClauq8OekC

Le Moullec, M., Pedersen, Å. Ø., Stien, A., Rosvold, J., & Hansen, B. B. (2019). A century of conservation: The ongoing recovery of Svalbard reindeer. In The Journal of Wildlife Management (Vol. 83, Issue 8, pp. 1676–1686). 10.1002/jwmg.21761

Leonardi, M., Librado, P., Der Sarkissian, C., Schubert, M., Alfarhan, A. H., Alquraishi, S. A., Al-Rasheid, K. A. S., Gamba, C., Willerslev, E., & Orlando, L. (2017). Evolutionary Patterns and Processes: Lessons from Ancient DNA. Systematic Biology, 66(1), e1– e29. 10.1093/sysbio/syw059

Li, H. (2013). Aligning sequence reads, clone sequences and assembly contigs with BWA- MEM. In arXiv *[q-bio.GN]*. arXiv. http://arxiv.org/abs/1303.3997

Li, H., Handsaker, B., Wysoker, A., Fennell, T., Ruan, J., Homer, N., Marth, G., Abecasis, G., Durbin, R., & 1000 Genome Project Data Processing Subgroup. (2009). The Sequence Alignment/Map format and SAMtools. Bioinformatics, 25(16), 2078–2079. 10.1093/bioinformatics/btp352

Lin, Z., Chen, L., Chen, X., Zhong, Y., Yang, Y., Xia, W., Liu, C., Zhu, W., Wang, H., Yan, B., Yang, Y., Liu, X., Kvie, K. S., Røed, K. H., Wang, K., Xiao, W., Wei, H., Li, G., Heller, R., … Li, Z. (2019). Biological adaptations in the Arctic cervid, the reindeer (Rangifer tarandus). Science, 364(6446). 10.1126/science.aav6312

Liu, D., Matzuk, M. M., Sung, W. K., Guo, Q., Wang, P., & Wolgemuth, D. J. (1998). Cyclin A1 is required for meiosis in the male mouse. Nature Genetics, 20(4), 377–380. 10.1038/3855

Li, Z., Lin, Z., Ba, H., Chen, L., Yang, Y., Wang, K., Qiu, Q., Wang, W., & Li, G. (2017). Draft genome of the reindeer (Rangifer tarandus). GigaScience, 6(12), 1–5. 10.1093/gigascience/gix102

Li, Z., Wang, L., Yue, H., Whitson, B. A., Haggard, E., Xu, X., & Ma, J. (2021). MG53, A Tissue Repair Protein with Broad Applications in Regenerative Medicine. Cells, 10(1). 10.3390/cells10010122

Loe, L. E., Liston, G. E., Pigeon, G., Barker, K., Horvitz, N., Stien, A., Forchhammer, M., Getz, W. M., Irvine, R. J., Lee, A., Movik, L. K., Mysterud, A., Pedersen, Å. Ø., Reinking, A. K., Ropstad, E., Trondrud, L. M., Tveraa, T., Veiberg, V., Hansen, B. B., & Albon, S. D. (2020). The neglected season: Warmer autumns counteract harsher winters and promote population growth in Arctic reindeer. Global Change Biology. 10.1111/gcb.15458

Lønø, O. (1959). REINEN PÅ SVALBARD. Norsk Polarinstitutt, Oslo, Norway. https://brage.npolar.no/npolar-xmlui/bitstream/handle/11250/2395010/Meddelelser083.pdf?sequence=1

Lorenzen, E. D., Nogués-Bravo, D., Orlando, L., Weinstock, J., Binladen, J., Marske, K. A., Ugan, A., Borregaard, M. K., Gilbert, M. T. P., Nielsen, R., Ho, S. Y. W., Goebel, T., Graf, K. E., Byers, D., Stenderup, J. T., Rasmussen, M., Campos, P. F., Leonard, J. A., Koepfli, K.-P., … Willerslev, E. (2011). Species-specific responses of Late Quaternary megafauna to climate and humans. Nature, 479(7373), 359–364. 10.1038/nature10574

Luypaert, T., Hagan, J. G., McCarthy, M. L., & Poti, M. (2020). Status of marine biodiversity in the Anthropocene. In YOUMARES 9-The Oceans: Our research, our future (pp. 57– 82). Springer, Cham. https://library.oapen.org/bitstream/handle/20.500.12657/22870/1007291.pdf?sequence=1#page=72

McKenna, A., Hanna, M., Banks, E., Sivachenko, A., Cibulskis, K., Kernytsky, A., Garimella, K., Altshuler, D., Gabriel, S., Daly, M., & DePristo, M. A. (2010). The Genome Analysis Toolkit: a MapReduce framework for analyzing next-generation DNA sequencing data. Genome Research, 20(9), 1297–1303. 10.1101/gr.107524.110

Meisner, J., & Albrechtsen, A. (2018). Inferring Population Structure and Admixture Proportions in Low-Depth NGS Data. Genetics, 210(2), 719–731. 10.1534/genetics.118.301336

Missler, M., Zhang, W., Rohlmann, A., Kattenstroth, G., Hammer, R. E., Gottmann, K., & Südhof, T. C. (2003). Alpha-neurexins couple Ca2+ channels to synaptic vesicle exocytosis. Nature, 423(6943), 939–948. 10.1038/nature01755

Mitchell, K. J., & Rawlence, N. J. (2021). Examining Natural History through the Lens of Palaeogenomics. Trends in Ecology & Evolution, 36(3), 258–267. 10.1016/j.tree.2020.10.005

Nei, M. (1987). Molecular Evolutionary Genetics. Columbia University Press. 10.7312/nei-92038

Nei, M., & Tajima, F. (1981). DNA polymorphism detectable by restriction endonucleases. Genetics, 97(1), 145–163. 10.1093/genetics/97.1.145

Nielsen, R., Korneliussen, T., Albrechtsen, A., Li, Y., & Wang, J. (2012). SNP calling, genotype calling, and sample allele frequency estimation from New-Generation Sequencing data. PloS One, 7(7), e37558. 10.1371/journal.pone.0037558

Paradis, E. (2010). pegas: an R package for population genetics with an integrated–modular approach. Bioinformatics, 26(3), 419–420. 10.1093/bioinformatics/btp696

Paradis, E., & Schliep, K. (2019). ape 5.0: an environment for modern phylogenetics and evolutionary analyses in R. Bioinformatics, 35(3), 526–528. 10.1093/bioinformatics/bty633

Peeters, B., Le Moullec, M., Raeymaekers, J. A. M., Marquez, J. F., Røed, K. H., Pedersen, Å. Ø., Veiberg, V., Loe, L. E., & Hansen, B. B. (2020). Sea ice loss increases genetic isolation in a high Arctic ungulate metapopulation. Global Change Biology, 26(4), 2028– 2041. 10.1111/gcb.14965

Peeters, B., Pedersen, Å. Ø., Loe, L. E., Isaksen, K., Veiberg, V., Stien, A., Kohler, J., Gallet, J.-C., Aanes, R., & Hansen, B. B. (2019). Spatiotemporal patterns of rain-on-snow and basal ice in high Arctic Svalbard: detection of a climate-cryosphere regime shift. Environmental Research Letters: ERL [Web Site], 14(1), 015002. 10.1088/1748-9326/aaefb3

Picard toolkit. (2019). In Broad Institute, GitHub repository. Broad Institute. http://broadinstitute.github.io/picard/

Pires-daSilva, A., Nayernia, K., Engel, W., Torres, M., Stoykova, A., Chowdhury, K., & Gruss, P. (2001). Mice deficient for spermatid perinuclear RNA-binding protein show neurologic, spermatogenic, and sperm morphological abnormalities. Developmental Biology, 233(2), 319–328. 10.1006/dbio.2001.0169

Puechmaille, S. J. (2016). The program structure does not reliably recover the correct population structure when sampling is uneven: subsampling and new estimators alleviate the problem. Molecular Ecology Resources, 16(3), 608–627. 10.1111/1755-0998.12512

Quinlan, A. R. (2014). BEDTools: The Swiss-Army Tool for Genome Feature Analysis. Current Protocols in Bioinformatics / Editoral Board, Andreas D. Baxevanis … [et Al.], 47, 11.12.1–34. 10.1002/0471250953.bi1112s47

R Core Team. (2021). R: A Language and Environment for Statistical Computing. R Foundation for Statistical Computing. https://www.R-project.org/

Reimer, P. J. (2020). Composition and consequences of the IntCal20 radiocarbon calibration curve. Quaternary Research, 96, 22–27. 10.1017/qua.2020.42

Reimers, E. (1984). Body composition and population regulation of Svalbard reindeer. Rangelands, 4(2), 16–21. 10.7557/2.4.2.499

Robin, M., Ferrari, G., Akgül, G., Münger, X., von Seth, J., Schuenemann, V. J., Dalén, L., & Grossen, C. (2022). Ancient mitochondrial and modern whole genomes unravel massive genetic diversity loss during near extinction of Alpine ibex. Molecular Ecology, 31(13), 3548–3565. 10.1111/mec.16503

Rohland, N., & Reich, D. (2012). Cost-effective, high-throughput DNA sequencing libraries for multiplexed target capture. Genome Research, 22(5), 939–946. 10.1101/gr.128124.111

Sánchez-Barreiro, F., Gopalakrishnan, S., Ramos-Madrigal, J., Westbury, M. V., de Manuel, M., Margaryan, A., Ciucani, M. M., Vieira, F. G., Patramanis, Y., Kalthoff, D. C., Timmons, Z., Sicheritz-Pontén, T., Dalén, L., Ryder, O. A., Zhang, G., Marquès-Bonet, T., Moodley, Y., & Gilbert, M. T. P. (2021). Historical population declines prompted significant genomic erosion in the northern and southern white rhinoceros (Ceratotherium simum). Molecular Ecology, 30(23), 6355–6369. 10.1111/mec.16043

Schubert, M., Ermini, L., Der Sarkissian, C., Jónsson, H., Ginolhac, A., Schaefer, R., Martin, M. D., Fernández, R., Kircher, M., McCue, M., Willerslev, E., & Orlando, L. (2014). Characterization of ancient and modern genomes by SNP detection and phylogenomic and metagenomic analysis using PALEOMIX. Nature Protocols, 9(5), 1056–1082. 10.1038/nprot.2014.063

Skotte, L., Korneliussen, T. S., & Albrechtsen, A. (2013). Estimating individual admixture proportions from next generation sequencing data. Genetics, 195(3), 693–702. 10.1534/genetics.113.154138

Song, R., Peng, W., Zhang, Y., Lv, F., Wu, H.-K., Guo, J., Cao, Y., Pi, Y., Zhang, X., Jin, L., Zhang, M., Jiang, P., Liu, F., Meng, S., Zhang, X., Jiang, P., Cao, C.-M., & Xiao, R.-P. (2013). Central role of E3 ubiquitin ligase MG53 in insulin resistance and metabolic disorders. Nature, 494(7437), 375–379. 10.1038/nature11834

Spielman, D., Brook, B. W., & Frankham, R. (2004). Most species are not driven to extinction before genetic factors impact them. Proceedings of the National Academy of Sciences of the United States of America, 101(42), 15261–15264. 10.1073/pnas.0403809101

Stempniewicz, L., Kulaszewicz, I., & Aars, J. (2021). Yes, they can: polar bears Ursus maritimus successfully hunt Svalbard reindeer Rangifer tarandus platyrhynchus. Polar Biology, 44(11), 2199–2206. 10.1007/s00300-021-02954-w

Taylor, R. S., Horn, R. L., Zhang, X., Golding, G. B., Manseau, M., & Wilson, P. J. (2019). The Caribou (Rangifer tarandus) Genome. Genes, 10(7), 540. 10.3390/genes10070540

Templeton, A. R., Crandall, K. A., & Sing, C. F. (1992). A cladistic analysis of phenotypic associations with haplotypes inferred from restriction endonuclease mapping and DNA sequence data. III. Cladogram estimation. Genetics, 132(2), 619–633. 10.1093/genetics/132.2.619

Teslovich, T. M., Musunuru, K., Smith, A. V., Edmondson, A. C., Stylianou, I. M., Koseki, M., Pirruccello, J. P., Ripatti, S., Chasman, D. I., Willer, C. J., Johansen, C. T., Fouchier, S. W., Isaacs, A., Peloso, G. M., Barbalic, M., Ricketts, S. L., Bis, J. C., Aulchenko, Y. S., Thorleifsson, G., … Kathiresan, S. (2010). Biological, clinical and population relevance of 95 loci for blood lipids. Nature, 466(7307), 707–713. 10.1038/nature09270

Therkildsen, N. O., Wilder, A. P., Conover, D. O., Munch, S. B., Baumann, H., & Palumbi, S. R. (2019). Contrasting genomic shifts underlie parallel phenotypic evolution in response to fishing. Science, 365(6452), 487–490. 10.1126/science.aaw7271

Trondrud, L. M., Pigeon, G., Król, E., Albon, S., Evans, A. L., Arnold, W., Hambly, C., Irvine, R. J., Ropstad, E., Stien, A., Veiberg, V., Speakman, J. R., & Loe, L. E. (2021). Fat storage influences fasting endurance more than body size in an ungulate. Functional Ecology, 35(7), 1470–1480. 10.1111/1365-2435.13816

Uhlén, M., Fagerberg, L., Hallström, B. M., Lindskog, C., Oksvold, P., Mardinoglu, A., Sivertsson, Å., Kampf, C., Sjöstedt, E., Asplund, A., Olsson, I., Edlund, K., Lundberg, E., Navani, S., Szigyarto, C. A.-K., Odeberg, J., Djureinovic, D., Takanen, J. O., Hober, S., … Pontén, F. (2015). Proteomics. Tissue-based map of the human proteome. Science, 347(6220), 1260419. 10.1126/science.1260419

UniProt Consortium. (2021). UniProt: the universal protein knowledgebase in 2021. Nucleic Acids Research, 49(D1), D480–D489. 10.1093/nar/gkaa1100

van der Bilt, W. G. M., D’Andrea, W. J., Werner, J. P., & Bakke, J. (2019). Early Holocene temperature oscillations exceed amplitude of observed and projected warming in Svalbard lakes. Geophysical Research Letters, 46(24), 14732–14741. 10.1029/2019gl084384

van der Knaap, W. O. (1989). Past Vegetation and Reindeer on Edgeoya (Spitsbergen) Between c. 7900 and c. 3800 BP, Studied by Means of Peat Layers and Reindeer Faecal Pellets. Journal of Biogeography, 16(4), 379–394. 10.2307/2845229

van der Valk, T., Díez-Del-Molino, D., Marques-Bonet, T., Guschanski, K., & Dalén, L. (2019). Historical Genomes Reveal the Genomic Consequences of Recent Population Decline in Eastern Gorillas. Current Biology: CB, 29(1), 165–170.e6. 10.1016/j.cub.2018.11.055

van Oort, B. E. H., Tyler, N. J. C., Gerkema, M. P., Folkow, L., Blix, A. S., & Stokkan, K.-A. (2005). Circadian organization in reindeer. Nature, 438(7071), 1095–1096. 10.1038/4381095a

Wang, W., Kissig, M., Rajakumari, S., Huang, L., Lim, H.-W., Won, K.-J., & Seale, P. (2014). Ebf2 is a selective marker of brown and beige adipogenic precursor cells. Proceedings of the National Academy of Sciences of the United States of America, 111(40), 14466– 14471. 10.1073/pnas.1412685111

Watterson, G. A. (1975). On the number of segregating sites in genetical models without recombination. Theoretical Population Biology, 7(2), 256–276. 10.1016/0040-5809(75)90020-9

Werner, K., Müller, J., Husum, K., Spielhagen, R. F., Kandiano, E. S., & Polyak, L. (2016). Holocene sea subsurface and surface water masses in the Fram Strait – Comparisons of temperature and sea-ice reconstructions. Quaternary Science Reviews, 147, 194–209. 10.1016/j.quascirev.2015.09.007

Williamsen, L., Pigeon, G., Mysterud, A., Stien, A., Forchhammer, M., & Loe, L. E. (2019). Keeping cool in the warming Arctic: thermoregulatory behaviour by Svalbard reindeer (Rangifer tarandus platyrhynchus). Canadian Journal of Zoology, 97(12), 1177–1185. 10.1139/cjz-2019-0090

Wollebaek, A. (1926). The Spitsbergen Reindeer:(rangifer Tarandus Spetsbergensis). Dybwad.

Yannic, G., Pellissier, L., Ortego, J., Lecomte, N., Couturier, S., Cuyler, C., Dussault, C., Hundertmark, K. J., Irvine, R. J., Jenkins, D. A., Kolpashikov, L., Mager, K., Musiani, M., Parker, K. L., Røed, K. H., Sipko, T., Þórisson, S. G., Weckworth, B. V., Guisan, A., … Côté, S. D. (2013). Genetic diversity in caribou linked to past and future climate change. Nature Climate Change, 4(2), 132–137. 10.1038/nclimate2074

Zhang, J., Bonasio, R., Strino, F., Kluger, Y., Holloway, J. K., Modzelewski, A. J., Cohen, P. E., & Reinberg, D. (2013). SFMBT1 functions with LSD1 to regulate expression of canonical histone genes and chromatin-related factors. Genes & Development, 27(7), 749–766. 10.1101/gad.210963.112

Zhang, K., Wang, T., Liu, X., Yuan, Q., Xiao, T., Yuan, X., Zhang, Y., Yuan, L., & Wang, Y. (2021). CASK, APBA1, and STXBP1 collaborate during insulin secretion. Molecular and Cellular Endocrinology, 520, 111076. 10.1016/j.mce.2020.111076

Zhang, Y., Wu, H.-K., Lv, F., & Xiao, R.-P. (2017). MG53: Biological Function and Potential as a Therapeutic Target. Molecular Pharmacology, 92(3), 211–218. 10.1124/mol.117.108241

