## Supporting Information for "A paleogenomic investigation of overharvest implications in an endemic wild reindeer subspecies"

### A paleo-genomic investigation of overharvest implications in an endemic wild reindeer subspecies

### Supplementary Information

##### **Table S1**. Comparison of mapping statistics between Svalbard reference genome and caribou reference genome. Values for each sample were generated as output after mapping with the PALEOMIX pipeline. This table is provided in a separate file.

##### **Table S2**. Summarized comparison of mapping statistics against different reference genomes, grouped by time frame. Tested reference assemblies were a consensus Svalbard reference genome, a caribou reference genome, and a reindeer mitochondrial genome

|  | Mean (min - max) sequencing depth of different reference genomes | | |
| --- | --- | --- | --- |
|  | **Svalbard reference** | **Caribou reference** | **Reindeer mitochondrial** |
| **Pre-hunting** | 1.42 (0.0004 -  6.78112) | 1.37 (0.0003 - 6.67) | 369.19 (30.11 - 1869.49) |
| **During hunting** | 4.44 (0.73 - 12.2) | 4.39 (0.72 - 12.14) | 218.77 (57.05 - 410.83) |
| **Ancient (pre+during)** | 2.37 (0.0004 - 12.22) | 2.32 (0.0003 - 12.14) | 340.16 (30.11 - 1869.49) |
| **Post-hunting** | 5.82 (0.02 - 58.89) | 5.74 (0.02 - 58.46) | 1484 (9.26 - 7907.88) |

**Table S3.** Metadata for all samples. This table is provided in a separate file**.**

##### **Table S4.** Identified proteins and genes in high-divergence windows. High divergence defined as high (>0.5) *F_ST_* as identified by sliding-window analysis. Proteomic information was retrieved from UniProt.

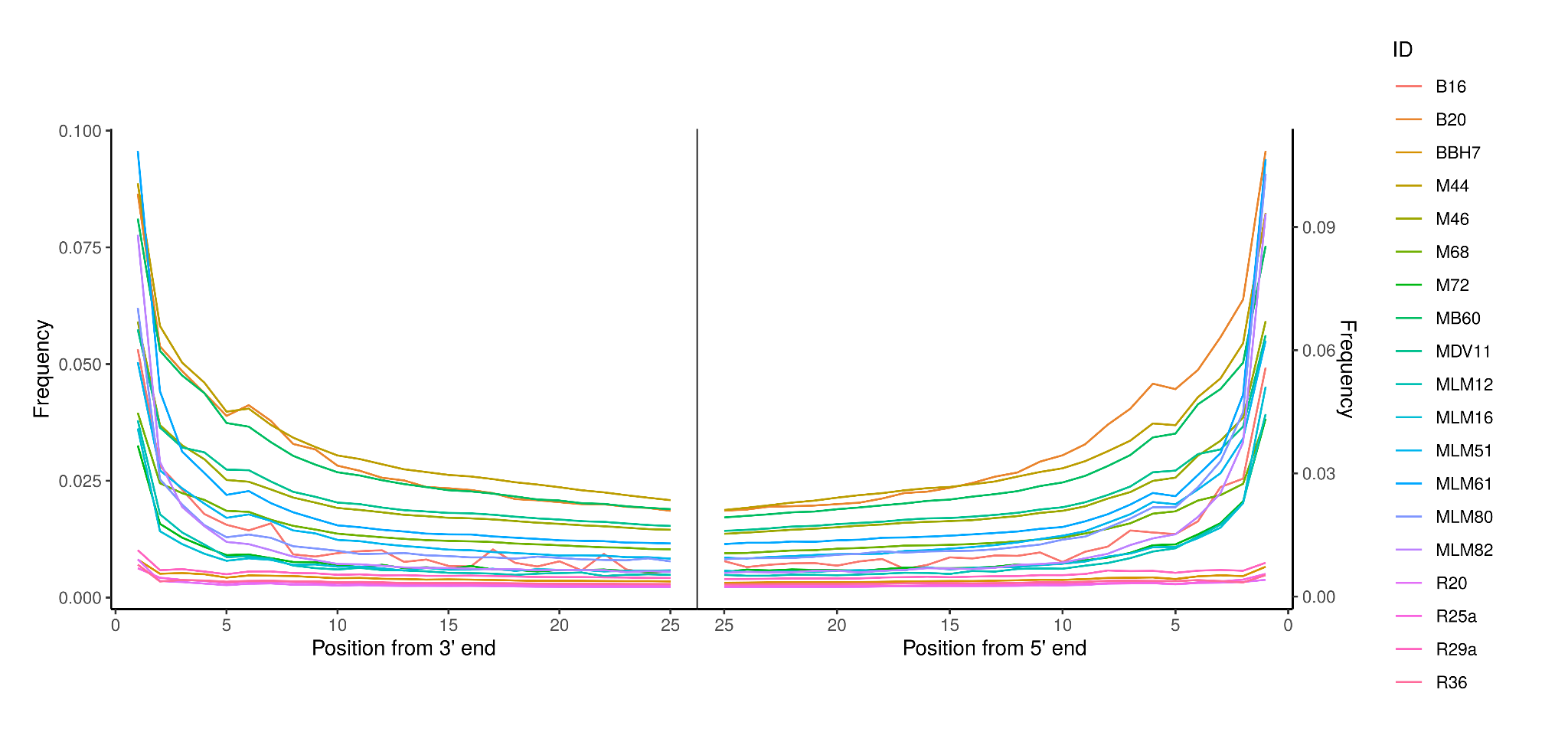

##### **Figure S5: DNA degradation patterns in ancient samples.** Frequency of base-misincorporation relative to the distance (in bases) from the 3’ end (left) or the 5’ end (right) of reads, respectively. Color denotes individual samples. Plotted values are from MapDamage2 output after clipped reads were removed.

##### **Figure S6:** **Population structure (admixture) barplot for K values between 2 and 10.** Each bar represents one individual Svalbard reindeer and the color represents affiliation to a proposed ancestral population. Individuals are grouped by spatiotemporal groups as defined by cluster analysis. NGSadmix was run 10 times for each K-value with all samples. Only the run with the highest likelihood is shown.

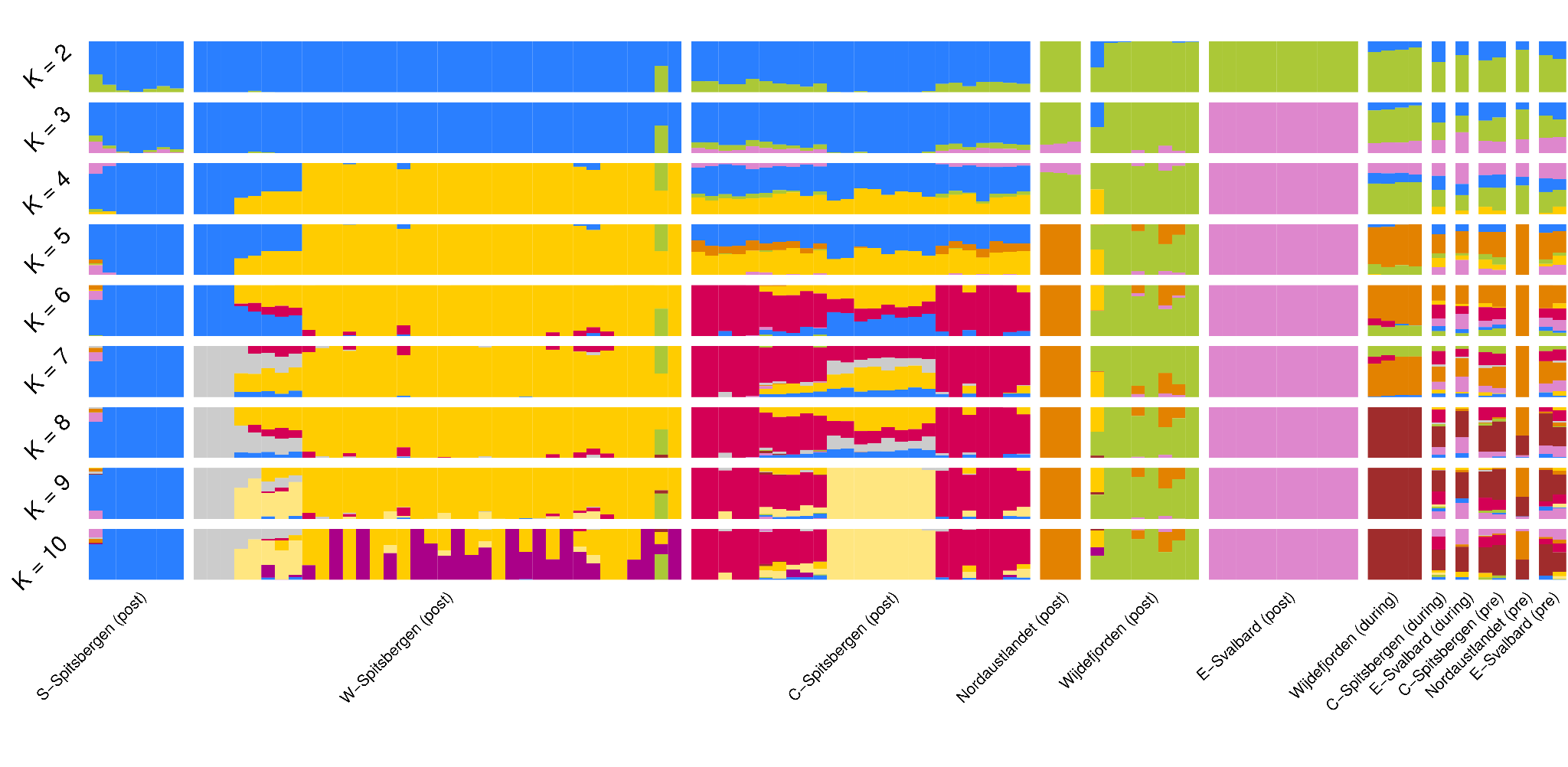

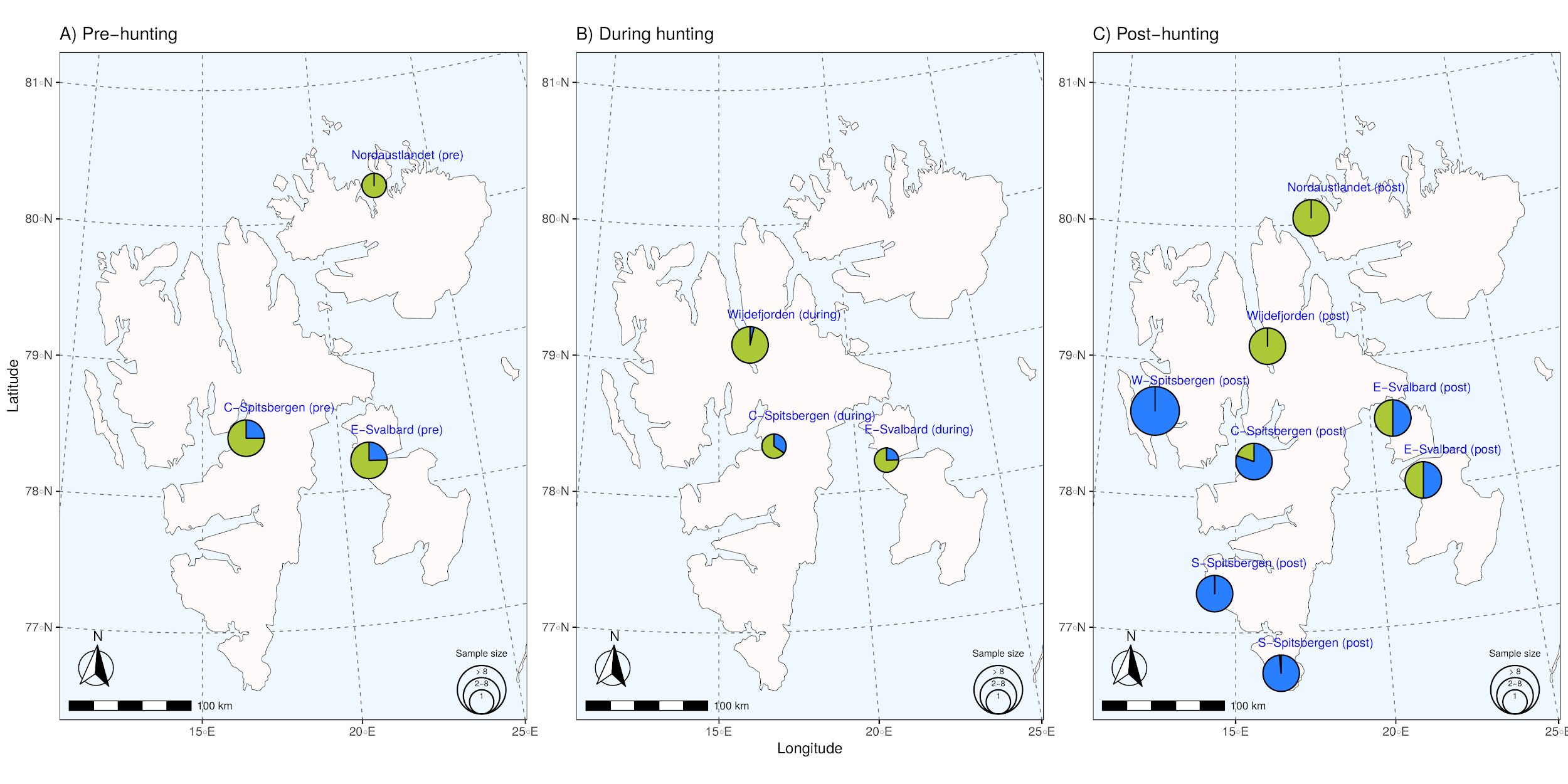

##### **Figure S7: Admixture proportions by sampling location for *K* = 2.** Individuals sampled within 80 km of each other are clustered together into the same pie. Pie size is scaled with the number of individuals. Individual ancestry proportions **B** before the hunting period, **C** during the hunting period, and **D** after the hunting period. C-Spitsbergen = Central Spitsbergen; S-Spitsbergen = South Spitsbergen; W-Spitsbergen = West Spitsbergen; E-Svalbard = East Svalbard.

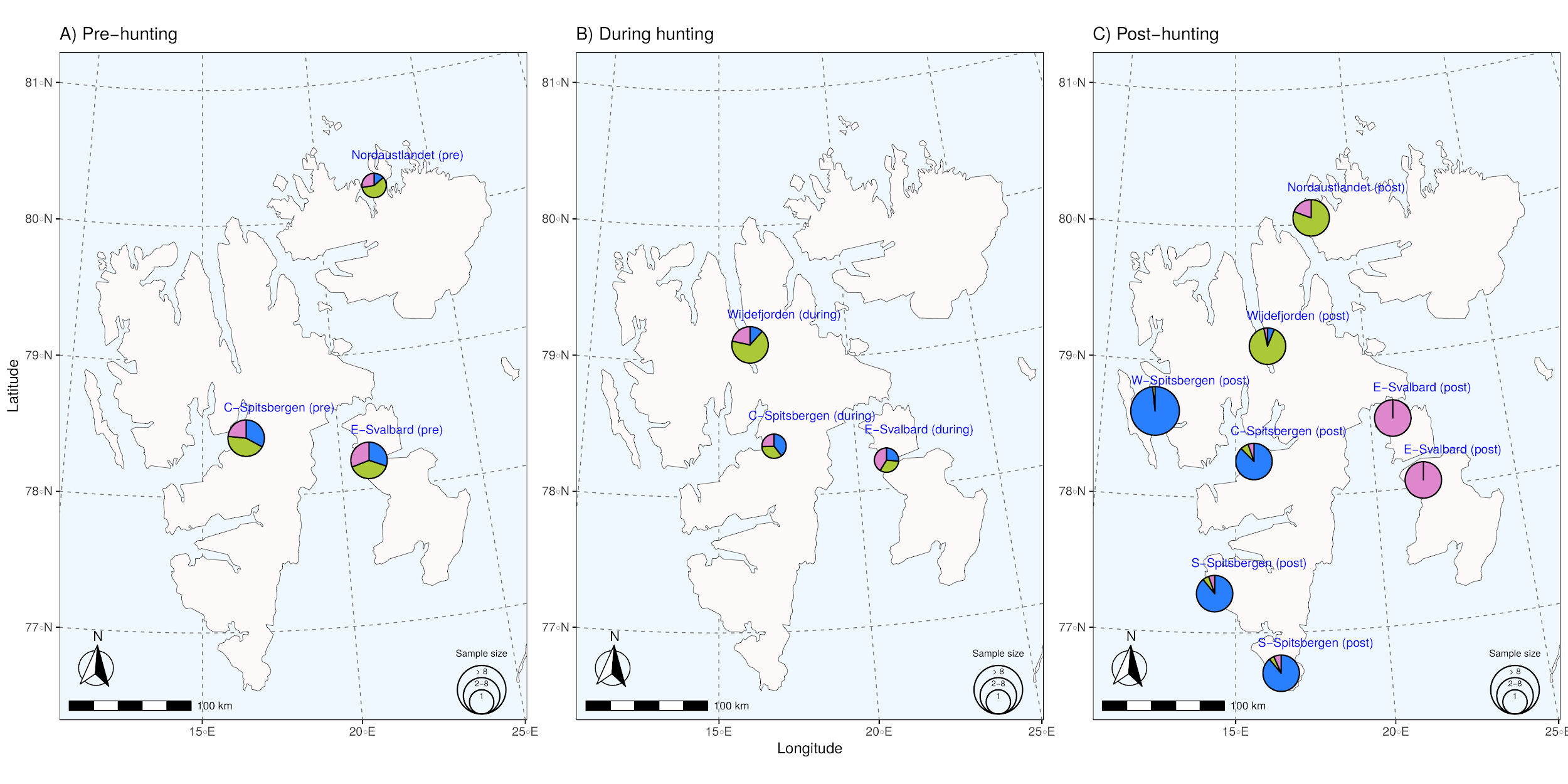

##### **Figure S8: Admixture proportions by sampling location for *K* = 3.** Individuals sampled within 80 km of each other are clustered together into the same pie. Pie size is scaled with the number of individuals. Individual ancestry proportions **B** before the hunting period, **C** during the hunting period, and **D** after the hunting period. C-Spitsbergen = Central Spitsbergen; S-Spitsbergen = South Spitsbergen; W-Spitsbergen = West Spitsbergen; E-Svalbard = East Svalbard.

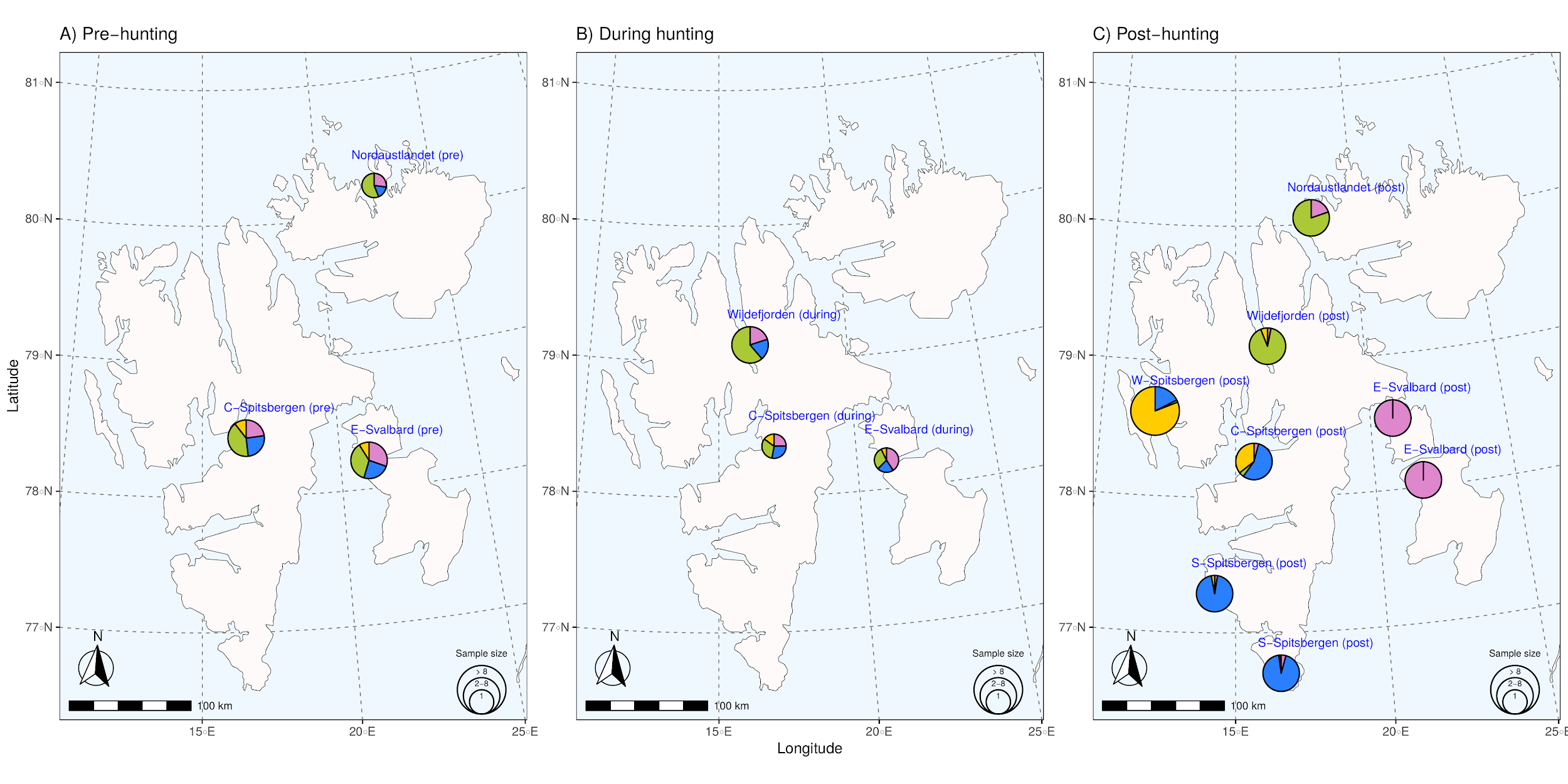

##### **Figure S9: Admixture proportions by sampling location for *K* = 4.** Individuals sampled within 80 km of each other are clustered together into the same pie. Pie size is scaled with the number of individuals. Individual ancestry proportions **B** before the hunting period, **C** during the hunting period, and **D** after the hunting period. C-Spitsbergen = Central Spitsbergen; S-Spitsbergen = South Spitsbergen; W-Spitsbergen = West Spitsbergen; E-Svalbard = East Svalbard.

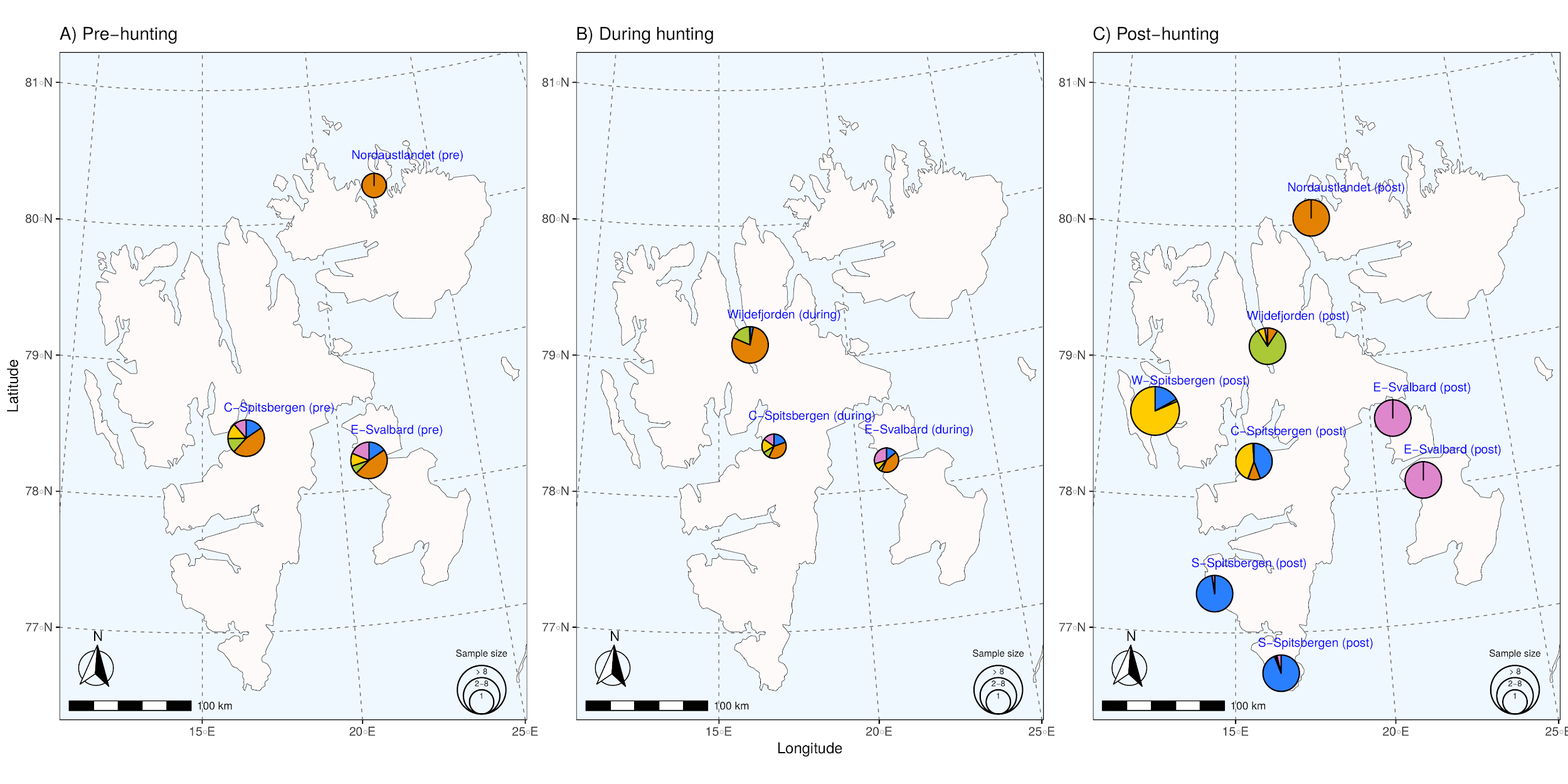

##### **Figure S10: Admixture proportions by sampling location for *K* = 5.** Individuals sampled within 80 km of each other are clustered together into the same pie. Pie size is scaled with the number of individuals. Individual ancestry proportions **B** before the hunting period, **C** during the hunting period, and **D** after the hunting period. C-Spitsbergen = Central Spitsbergen; S-Spitsbergen = South Spitsbergen; W-Spitsbergen = West Spitsbergen; E-Svalbard = East Svalbard.

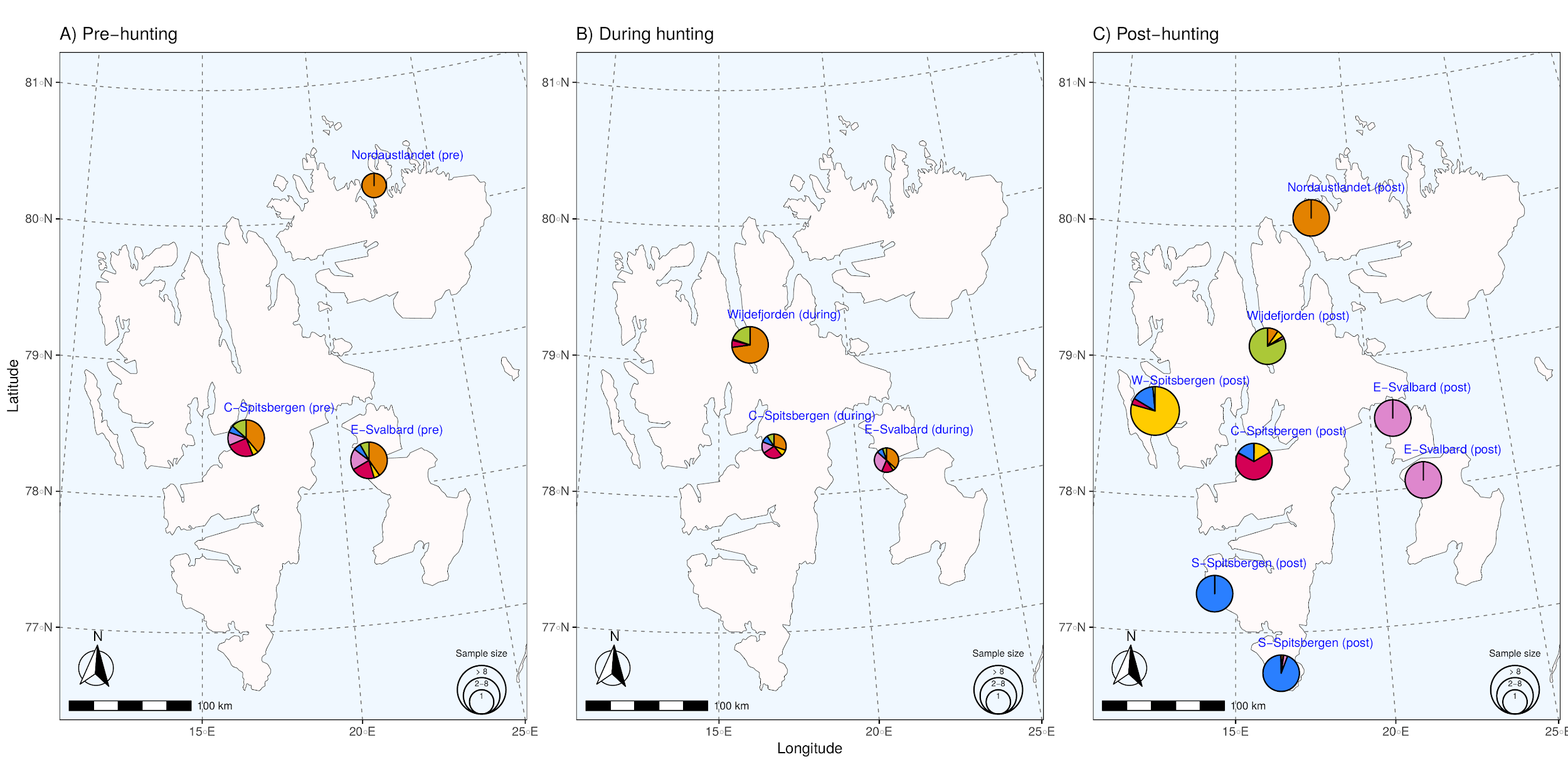

##### **Figure S11: Admixture proportions by sampling location for *K* = 6.** Individuals sampled within 80 km of each other are clustered together into the same pie. Pie size is scaled with the number of individuals. Individual ancestry proportions **B** before the hunting period, **C** during the hunting period, and **D** after the hunting period. C-Spitsbergen = Central Spitsbergen; S-Spitsbergen = South Spitsbergen; W-Spitsbergen = West Spitsbergen; E-Svalbard = East Svalbard.

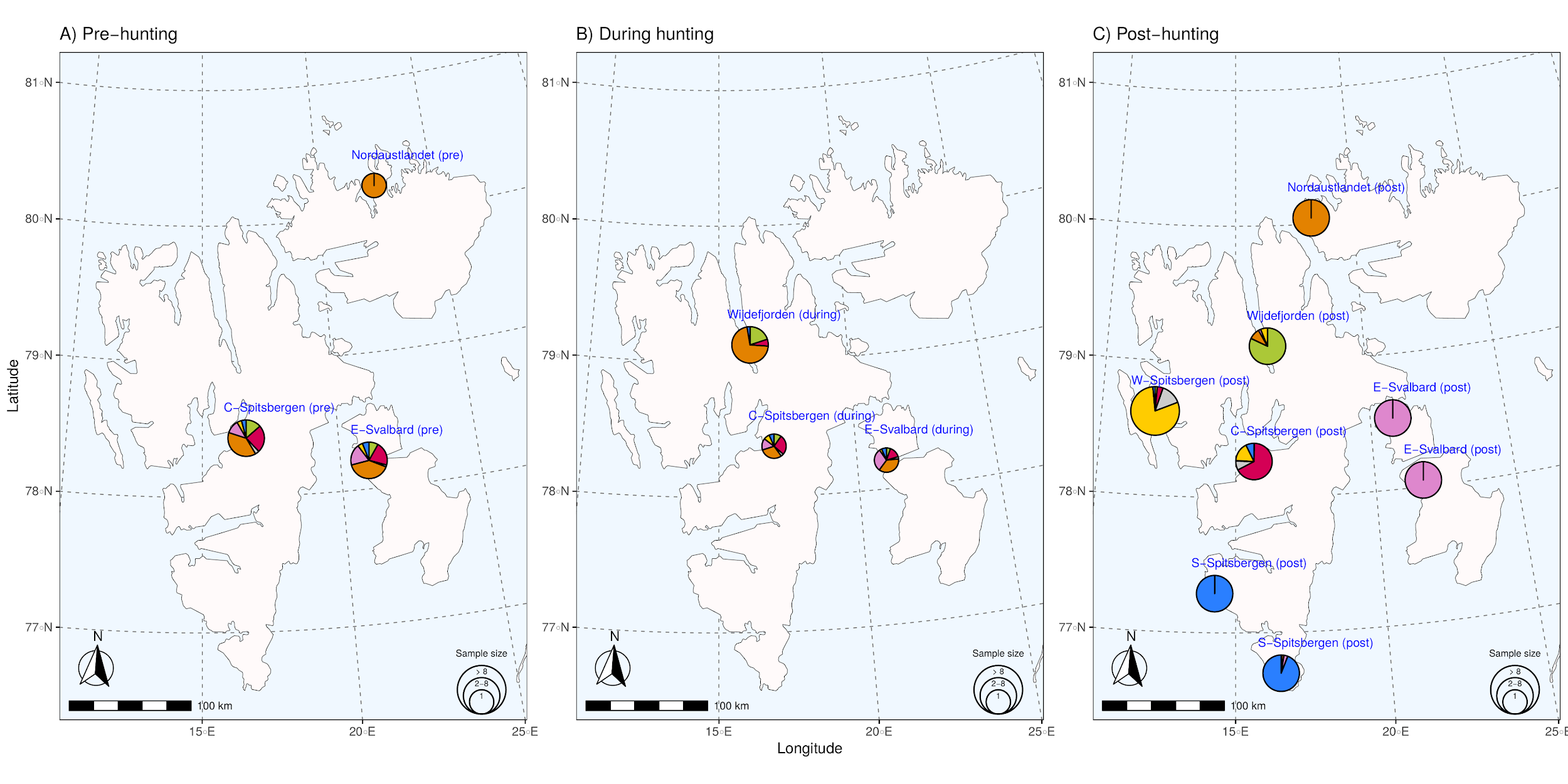

##### **Figure S12: Admixture proportions by sampling location for *K* = 7.** Individuals sampled within 80 km of each other are clustered together into the same pie. Pie size is scaled with the number of individuals. Individual ancestry proportions **B** before the hunting period, **C** during the hunting period, and **D** after the hunting period. C-Spitsbergen = Central Spitsbergen; S-Spitsbergen = South Spitsbergen; W-Spitsbergen = West Spitsbergen; E-Svalbard = East Svalbard.

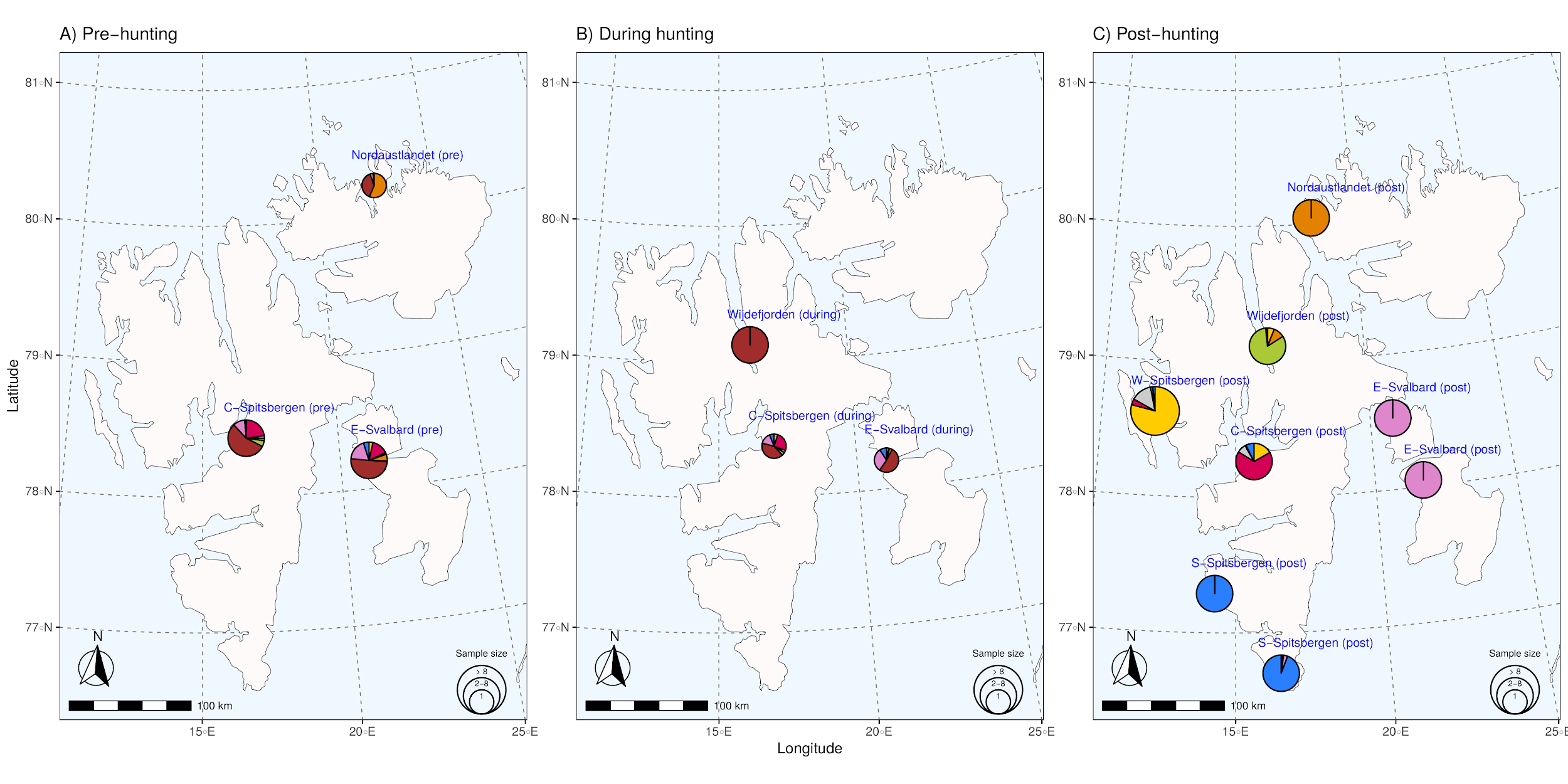

##### **Figure S13: Admixture proportions by sampling location for *K* = 8.** Individuals sampled within 80 km of each other are clustered together into the same pie. Pie size is scaled with the number of individuals. Individual ancestry proportions **B** before the hunting period, **C** during the hunting period, and **D** after the hunting period. C-Spitsbergen = Central Spitsbergen; S-Spitsbergen = South Spitsbergen; W-Spitsbergen = West Spitsbergen; E-Svalbard = East Svalbard.

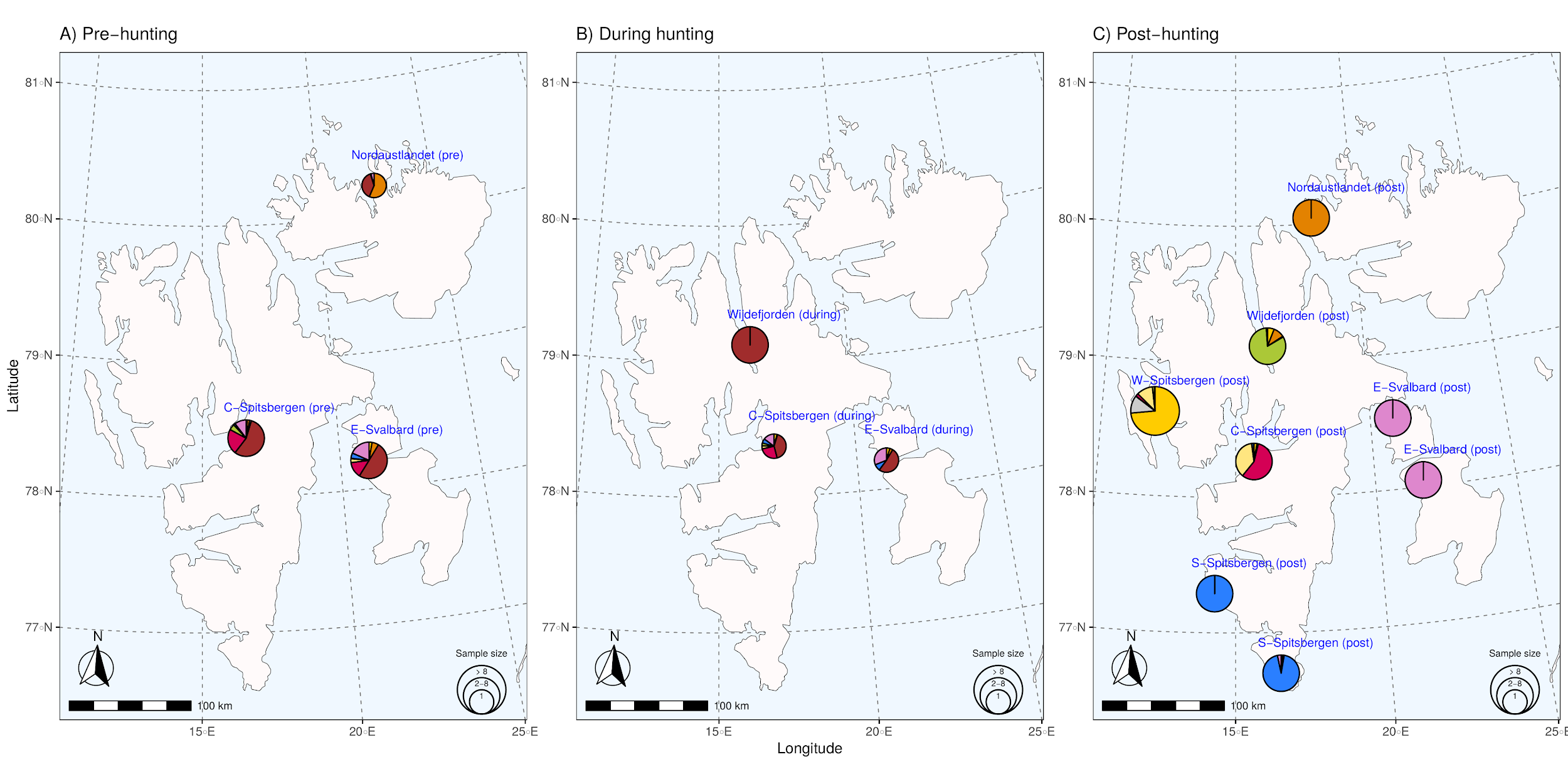

##### **Figure S14: Admixture proportions by sampling location for *K* = 9.** Individuals sampled within 80 km of each other are clustered together into the same pie. Pie size is scaled with the number of individuals. Individual ancestry proportions **B** before the hunting period, **C** during the hunting period, and **D** after the hunting period. C-Spitsbergen = Central Spitsbergen; S-Spitsbergen = South Spitsbergen; W-Spitsbergen = West Spitsbergen; E-Svalbard = East Svalbard.

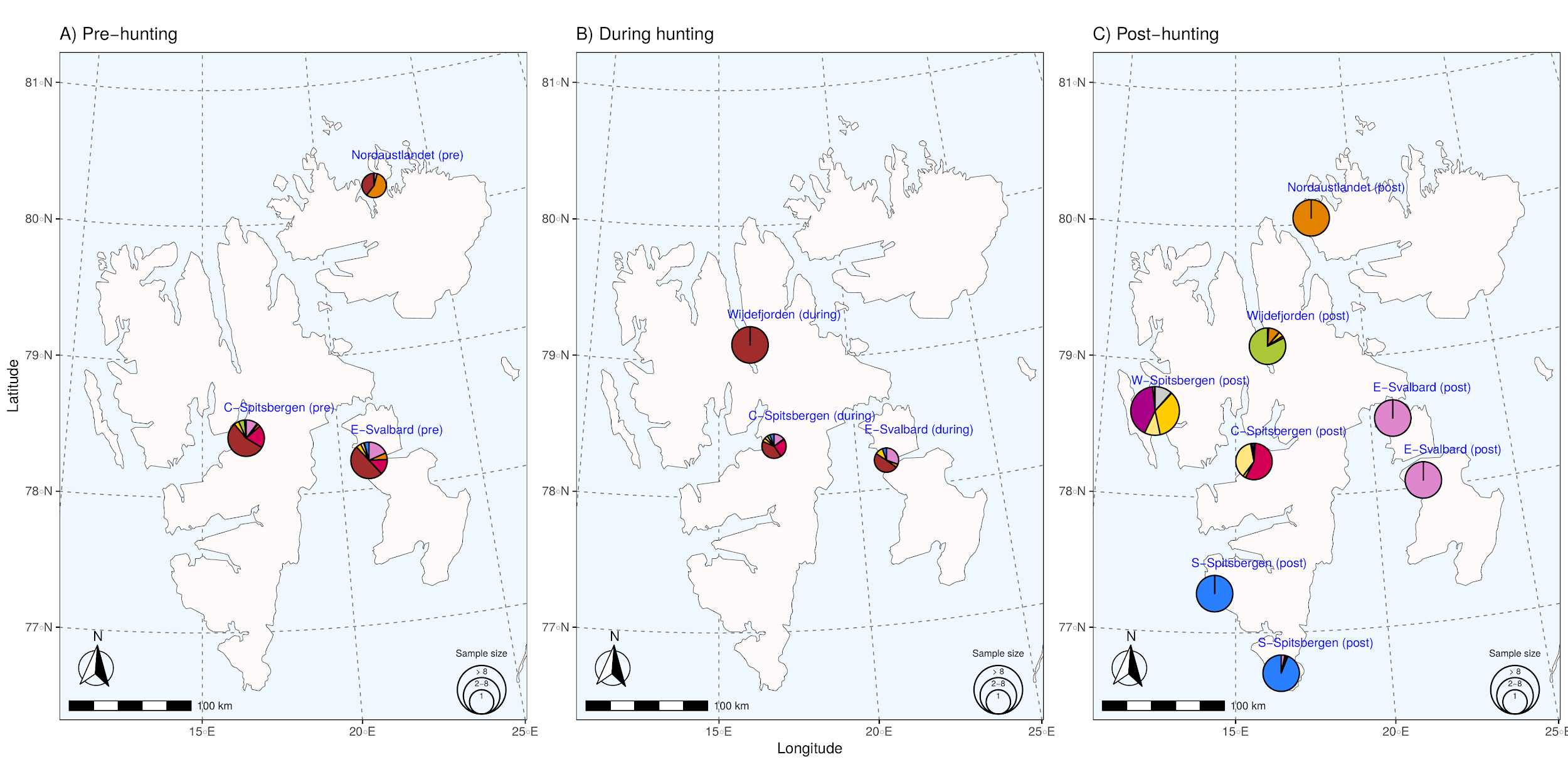

##### **Figure S15: Admixture proportions by sampling location for *K* = 10.** Individuals sampled within 80 km of each other are clustered together into the same pie. Pie size is scaled with the number of individuals. Individual ancestry proportions **B** before the hunting period, **C** during the hunting period, and **D** after the hunting period. C-Spitsbergen = Central Spitsbergen; S-Spitsbergen = South Spitsbergen; W-Spitsbergen = West Spitsbergen; E-Svalbard = East Svalbard.

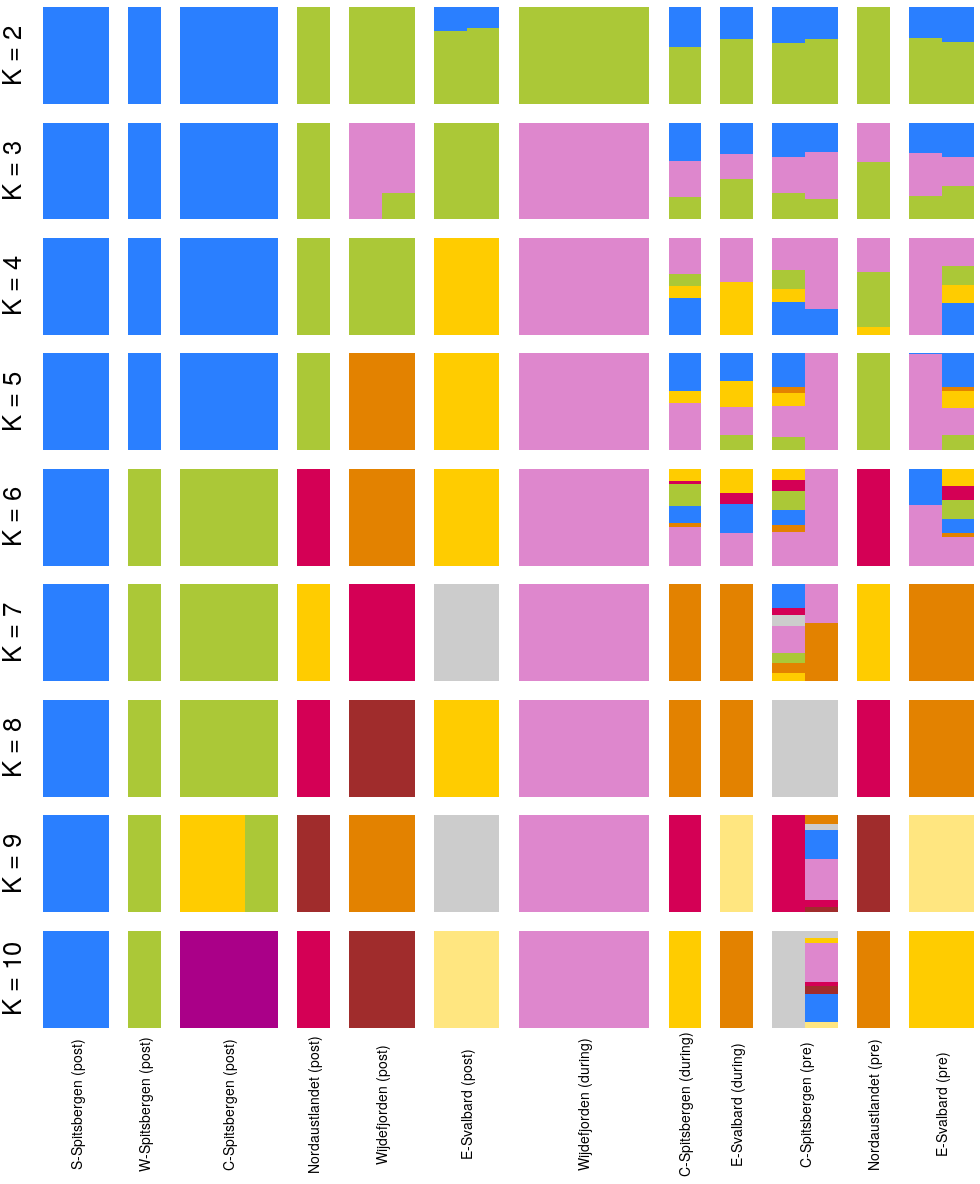

##### **Figure S16:** **Population structure (admixture) barplot for K values between 2 and 10 and a dataset of reduced modern sample size.** Each bar represents one individual Svalbard reindeer and the color represents affiliation to a proposed ancestral population. Individuals are grouped by spatiotemporal groups as defined by cluster analysis. NGSadmix was run 10 times for each K-value with all samples. Only the run with the highest likelihood is shown.

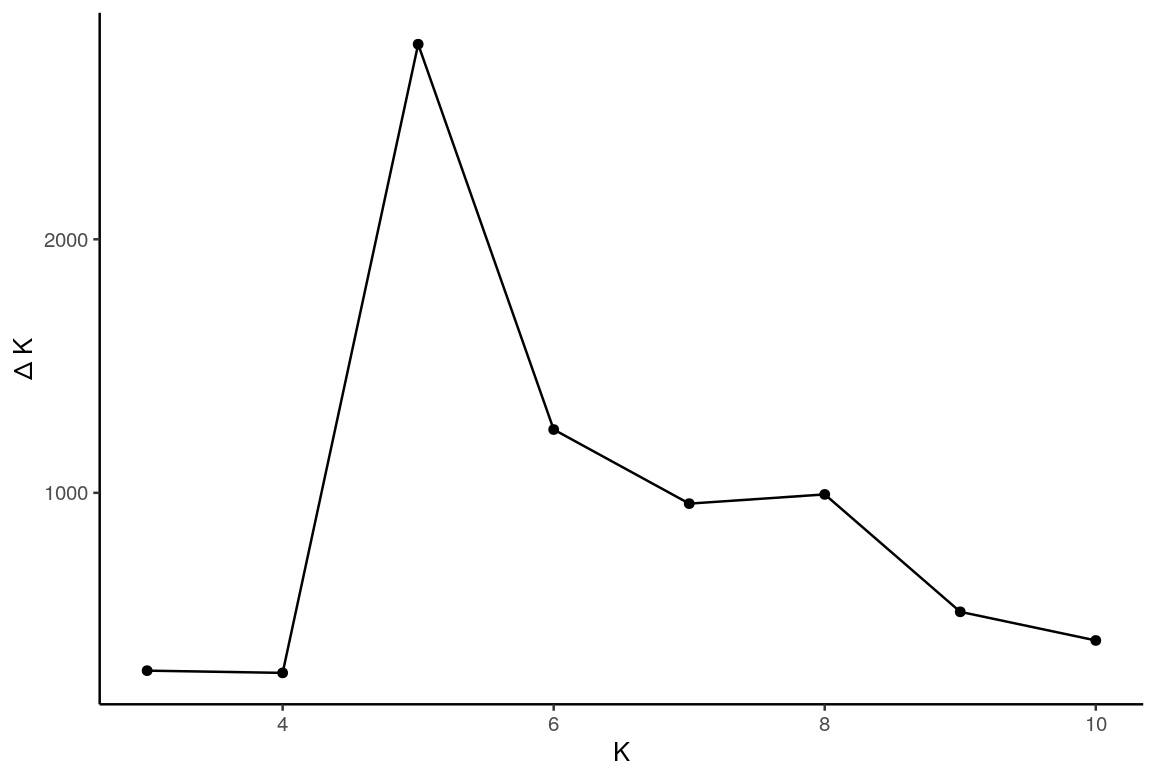

##### **Figure S17: Delta-K graph used to estimate the optimal value of *K*.**

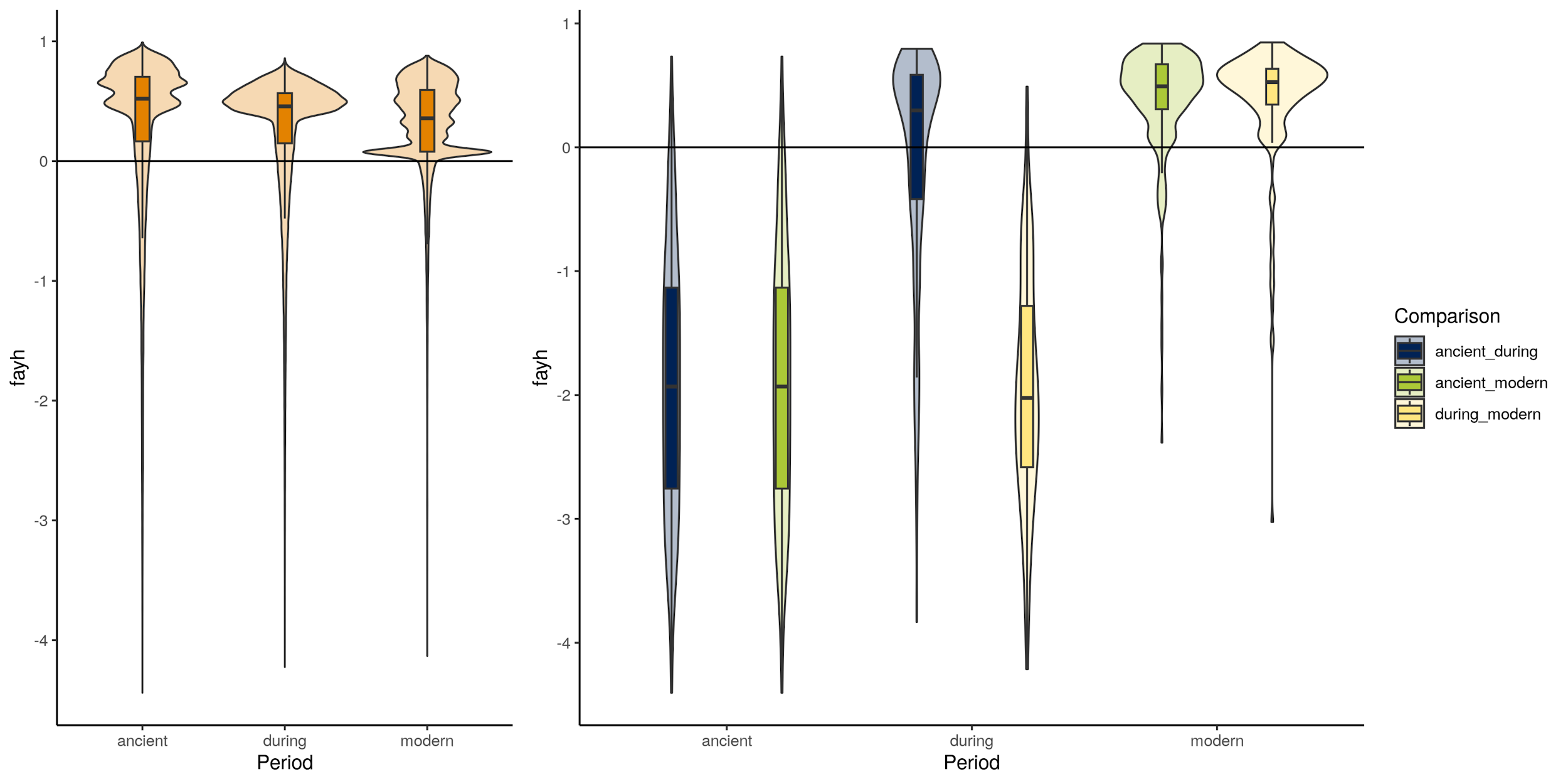

##### **Figure S18: Fay & Wu’s H.** Fay & Wu’s H for samples from the during-hunting and post-hunting period, respectively outlier and non-outlier windows. Bottom left: F_ST_ non-outlier windows. Bottom right: F_ST_ outlier windows.
